# Unifying physical and molecular coordinate systems across modalities in spatial biology

**DOI:** 10.64898/2026.09.01.748536

**Authors:** Bingjie Dai, Yunzhi Yan, Zhikang Wang, Yuchao Liang, Sijie Li, Pengwei Hu, Xinwang Yang, Chuyao Wang, Litai Yi, Chenglong Sun, Jinhai Huang, Xingtao Zhou, Hebing Chen, Daoliang Zhang, Qi Zou, Yixuan Du, Zheqi Hu, Yongqiang Xing, Guifang Cao, Zhenxing Feng, Jianfeng Feng, Shuhua Xu, Wei Hu, Yongchun Zuo, Bin-Zhi Qian, Zhiyuan Yuan

## Abstract

Establishing a unified physical and molecular coordinate system from fragmented multi-modal data is a longstanding challenge in biology. Here, we present MAPS, a modality-agnostic platform for spatial biology comprising (1) MAPS-alignment for ultrafast alignment of any modality, (2) MAPS-integration for both anchored and unanchored integration across orthogonal modalities for 3D multi-modal reconstruction, and (3) MAPS-Explorer for large-scale interactive 3D analysis. MAPS outperformed existing methods across extensive benchmarks on 34 datasets spanning 16 technology platforms and 6 modalities, while delineating fine-grained multi-modal tissue architectures across diverse biological systems in mouse and human. At cross-consortium scale, MAPS integrated 434 slices comprising 21 million cells from 18 atlases and 5 modalities to construct the most comprehensive 3D multi-modal mouse brain atlas. At individual laboratory scale, MAPS empowered routine 2D spatial assays to reconstruct continuous 3D multi-modal landscapes of human hepatocellular carcinoma, revealing the limitations of 2D spatial relationships and uncovering depth-dependent immune-state transitions.

## Introduction

The structural and functional principles of multicellular life are governed by the interplay between high-dimensional cellular molecular states and their physical spatial coordinates within the native tissue architecture^1,2^. Spatially resolved profiling technologies have shifted the paradigm of cellular cartography, enabling the construction of cellular coordinate systems directly from *in situ* spatial data rather than from dissociated single-cell measurements^3,4^. However, current spatial measurements remain fundamentally fragmented, largely constrained to single modalities and thin 2D tissue slices^4-31^. Consequently, the field still lacks a unified coordinate framework that links physical structure to molecular state across sections and modalities, thereby enabling comprehensive 3D reconstruction of multicellular systems.

The need for constructing such unified physical and feature coordinate systems spans a broad spectrum of spatial biology applications, exemplified by two prominent paradigms at either end of the spectrum. At one end of the spectrum, large-scale atlas-building consortia^32-36^ require harmonizing massive, heterogeneous datasets across diverse spatial modalities, platforms, and laboratories, demanding a highly scalable and modality-agnostic spatial alignment engine. At the other end, individual research and translational teams increasingly seek accessible workflows to construct bespoke 3D coordinate systems de novo for specific specimens, such as serial slices of patient tumor biopsies^37^. This typically entails registering consecutive slices and bridging orthogonal feature spaces across distinct assays from the same tissue block. Across these archetypes and intermediate use cases, there is an overarching need for a computational platform characterized by high scalability, modality and omics agnosticism, versatile support for both anchored and unanchored multi-modal spatial data, and interactive 3D multi-modal visualization of the resulting reconstructions at the scale of tens of millions of cells.

Despite the foundational importance of such infrastructure, current computational methods tackle physical space registration and molecular state integration in isolation, leaving the full spectrum of requirements unmet. On the one hand, physical space alignment methods, such as PASTE^38^, Spateo^39^, CAST^40^, 3d-OT^41^, and GPSA^42^, predominantly rely on shared molecular features or are tailored to specific modalities to guide coordinate transformation. Consequently, such methods fail to accommodate orthogonal modalities with non-overlapping feature spaces, and also suffer from prohibitive computational overhead on modern datasets, as highlighted in recent benchmarks^43^. On the other hand, feature space integration methods are typically confined to single-omics profiles (e.g., NicheCompass^44^ and FuseMap^45^) or strictly paired, co-assayed measurements (e.g., MISO^46^ and SpatialGlue^47^), whereas methods that rely on intermediate shared modality often struggle to simultaneously reconcile complex mixtures of anchored and un anchored datasets^48,49^. Critically, no existing framework unifies physical alignment and feature integration into a modality-agnostic, scalable, and cohesive pipeline, nor does the community possess an interactive visualization infrastructure capable of seamless exploration and interrogation of the resulting large-scale, multi-modal physical and state coordinate systems.

Here, we introduce MAPS (Modality-Agnostic alignment Platform for Spatial biology), a comprehensive computational framework designed to construct unified physical coordinate systems and integrate multi-omics feature spaces across arbitrary modalities. The MAPS platform comprises three core pillars: (1) MAPS-alignment, a modality-agnostic and ultrafast physical alignment module that operates purely on geometry, eliminating reliance on shared cellular features and resolving orthogonal cross-modal alignment across diverse platforms; (2) MAPS-integration, comprising MAPS-integration-U and MAPS-integration-P, a versatile feature integration engine tailored to distinct experimental paradigms. MAPS-integration-U performs unanchored diagonal integration across spatially continuous or adjacent slices by propagating orthogonal modal features across a physically aligned coordinate system; MAPS-integration-P empowers accessible, cost-effective laboratory workflows by utilizing available modalities as structural anchors to guide 3D multi-modal integration across tissue blocks with substantial morphological variation; (3)MAPS-Explorer, a cross-platform, high-performance interactive visualization platform capable of rendering tens of millions of cells and supporting diverse multi-dimensional analyses, enabling both cross-atlas exploration and specimen-specific discovery.

We systematically evaluated MAPS across 34 benchmark datasets spanning 16 technology platforms and 6 modalities (Supplementary Fig. 2), including spatial epigenomics^27-31^, transcriptomics^6-17^, translatomics^18,19^, proteomics^24-26^, metabolomics^20-23^ and high-resolution tissue images, such as Hematoxylin and Eosin (H&E) and Immunofluorescence (IF) images, demonstrating robust invariance of MAPS to rotations, coordinate offsets, scaling factors, and partial tissue overlap. MAPS-alignment outperformed 7 state-of-the-art alignment methods in 5 types of intra-omics alignment scenarios, achieving an average speed up of more than 100-fold and a substantial reduction in computational resource consumption. For example, it aligned two breast cancer slices containing ∼900,000 cells each in 41 s, with a peak GPU memory usage of 4.8 GB. Notably, MAPS-alignment performed single-cell-level alignment of mouse embryo and H&E images (containing >1.35 million cells) within 76 s, achieving accuracy comparable to the manual proofreading of Xenium official, while requiring less than 5 GB of GPU memory. Beyond alignment, MAPS-integration establishes a unified framework for multi-modal analysis across both anchored and un anchored settings by synergistically coupling morphology with cross-modal associations. In unanchored scenarios involving arbitrary orthogonal modalities without shared features, MAPS-integration is, to our knowledge, the only method capable of such integration. It accurately resolves the triple-modal anatomical architecture of the mouse cerebellum and uncovers functional regionalization in the mouse tongue epithelium. In anchored settings, using paired H&E histology as the anchor to predict spatial multi-omics at cellular resolution, MAPS-integration outperformed seven state-of-the-art omics prediction methods, with Pearson correlation coefficients improving by 30%-50%.

We further demonstrate that MAPS supports the full spectrum of spatial biology applications, from large-scale atlas-building consortia to individual research and translational teams. To demonstrate its utility at the cross-consortium end of the spectrum, we applied MAPS to assemble 434 slices comprising over 21 million cells from 18 atlases, 14 platforms, and 5 modalities, establishing the most comprehensive multi-modal mouse brain atlas in the field. Meanwhile, at the accessible individual-laboratory end, we applied MAPS to an in-house series of slices (containing ∼26 million cells) from a human hepatocellular carcinoma (HCC) specimen, anchored by dense serial H&E imaging and sparse multi-omics profiling, thus reconstructing a continuous 3D spatial multi-omics landscape. 3D distance measurements uncovered systematic underestimation of cellular proximity in 2D representations, leading to skewed interpretations of cellular neighborhood composition and potential cell interactions. Beyond revealing spatial biases, MAPS revealed depth-dependent immune-state transitions, spatially structured monocyte distributions.

## Result

### Overview of MAPS

MAPS constitutes a comprehensive computational platform for aligning and integrating diverse spatial modalities, including epigenomics, transcriptomics, translatomics, proteomics, metabolomics and imaging data (e.g., H&E and IF) (Fig. 1a). MAPS can establish a unified physical coordinate system and integrate multi-omics representation spaces across arbitrary spatial platforms (Fig. 1b). The MAPS framework comprises three core components: MAPS-alignment (Supplementary Fig 1a; see methods) for modality-agnostic, ultra-fast, geometry-driven physical alignment, MAPS-integration for versatile multi-modal feature integration (Supplementary Fig. 1b, c; see methods), and MAPS-Explorer for real-time interactive 3D exploration and spatial analytics (Fig.1e, f; see methods).

**Fig. 1.**
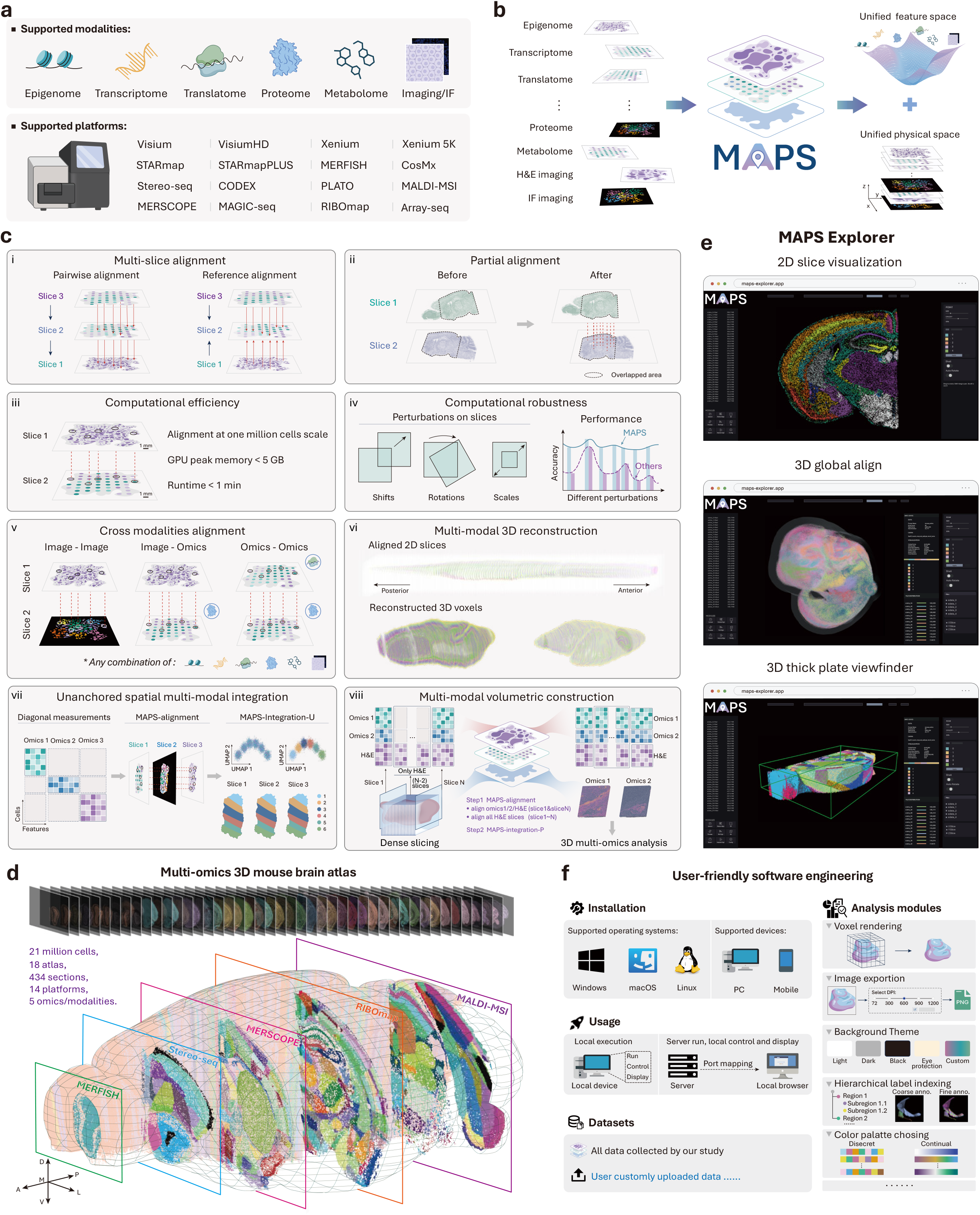
Overview of MAPS. **a**, MAPS enables alignment and integration across diverse spatial modalities, including spatial omics data and H&E images. **b**, MAPS unifies physical and `feature space` across modalities for spatial biology. **c**, Core characteristics of MAPS. MAPS provides two multi-slice alignment strategies: pairwise alignment (i), which iteratively aligns adjacent slices, and reference alignment, which aligns all slices to a common reference slice. It also supports partial alignment for slices with only partially overlapping regions (ii). MAPS is computationally efficient, aligning one million spots in 1 minute with less than 5 GB GPU memory (iii). MAPS is computationally robust against induced perturbations (iv). In addition, MAPS enables cross-modal alignment between arbitrary spatial modalities (v), enabling multi-modal 3D reconstruction (vi). Furthermore, MAPS-integration enables unpaired cross-modal integration across orthogonal modalities (vii) and histology-anchored multi-modal volumetric construction (viii). **d**, A multi-modal 3D mouse brain atlas constructed using MAPS. The atlas integrates datasets from 5 spatial modalities and 14 technology platforms. **e**, Overview of MAPS-Explorer. MAPS-Explorer provides an interactive interface for result visualization and analysis. **f**, Overview of the MAPS software ecosystem. MAPS offers multiple installation and deployment options, and integrates result exploration for this study, user data upload, and a collection of downstream analysis modules.

To establish precise spatial correspondence across disparate modalities without relying on shared molecular features, MAPS-alignment operates directly on spatial geometry (Supplementary Fig. 1a). MAPS-alignment supports diverse experimental setups. For multi-slice alignment (Fig. 1c (i)), MAPS-alignment supports two modes. Pairwise alignment sequentially aligns neighboring slices for serial tracking, whereas reference alignment aligns all slices to a designated reference slice. In addition, partial alignment is specifically designed to handle slices with incomplete spatial overlap of tissue (Fig. 1c (ii)). Owing to its lightweight algorithmic architecture (Supplementary Fig 1a), MAPS-alignment scales efficiently to large datasets, processing one million cells in less than one minute with peak GPU memory usage below 5 GB (Fig. 1c (iii) and Supplementary Fig. 2 last two columns). Furthermore, MAPS-alignment maintains robust performance under coordinate perturbations, including shifts, rotations, and scaling (Fig. 1c (iv)), establishing the first applicable alignment solution capable of modality-agnostic (Fig. 1c (v)) and 3D spatial reconstruction across diverse systems (Fig. 1c (vi)).

Building upon precisely aligned physical coordinates, the MAPS-integration module executes downstream feature integration through two complementary integration engines tailored to distinct experimental paradigms (Fig. 1c (vii, viii)). For spatially contiguous or adjacent slices profiled across non-overlapping omics layers, MAPS-integration-U performs unanchored integration by computationally propagating unmeasured molecular states across physically aligned continuous spaces (Fig. 1c (vii) and Supplementary Fig. 1b). Conversely, for tissue blocks exhibiting substantial morphological variation or multi-batch distortions, MAPS-integration-P provides an accessible and cost-effective workflow by utilizing readily accessible and low-cost modalities (e.g., histology) as structural anchors to guide paired 3D multi-modal integration and cross-slice molecular imputation (Fig. 1c (viii) and Supplementary Fig. 1c).

We applied MAPS-alignment to construct a cross-consortium multi-modal 3D mouse brain atlas in spatial biology, encompassing 434 slices with over 21 million cells derived from 18 atlases, 14 platforms, and 5 modalities (Fig. 1d). To facilitate seamless exploration, visual interrogation, and data sharing of such massive volumetric multi-modal datasets, we developed MAPS-Explorer, an open-source, high-performance interactive 3D visualization and analytical platform (Fig. 1e). MAPS-Explorer supports cross-platform and cross-device deployment, serverless read-only web deployment for public dissemination, and smooth real-time rendering of atlas-scale datasets exceeding containing tens of millions of cells. Beyond visualization, MAPS-Explorer also provided diverse analysis functional, including dynamic information statistics, 2D/3D visualization, 3D model animation, cell density viewer, thick plate viewing, voxel rendering, background theme, hierarchical label indexing, palette selection, 3D surfaces dynamics analysis, multi gene co-expression visualization, 3D cells communication visualization, custom parameter panel, and adjustable resolution image export, among others (Fig. 1f).

### MAPS-alignment delivers accurate, robust, ultrafast, and scalable alignment of intra-modal spatial data

Reconstructing 3D tissue architecture from serial 2D slices within the same molecular modality requires precise spatial alignment into a unified coordinate system. In experimental workflows, serial slices can acquire rotational, translational and field-of-view discrepancies during sectioning, mounting and imaging. Furthermore, spatial alignment is hindered by measurement noise, variable capture efficiencies, and genuine biological variation across serial slices. While numerous spatial alignment algorithms have been developed^42,50-55^, most of them were designed primarily for spatial transcriptomics (ST). Recent systematic benchmarks^43^ have further shown that existing tools frequently suffer from severe computational bottlenecks such as long runtimes, and out-of-memory (OOM) failures on large-scale datasets, and an inability to handle cross-platform alignment. Therefore, we reasoned that a robust, geometry-based alignment engine should first establish exceptional accuracy, efficiency, and scalability in standard and challenging intra-modal tasks before extending to inter-modal paradigms.

To systematically benchmark MAPS-alignment against 7 state-of-the-art methods^42,50-55^, we assembled a comprehensive collection of spatial transcriptomics datasets generated across ten distinct technological platforms, including 10x Visium, Visium HD, Xenium, MERFISH, STARmap PLUS, Stereo-seq, ST, CosMx, Array-seq and Slide-seq, as well as diverse biological samples, such as human breast cancer^9^, human brain^56^, mouse brain^57^, mouse and non-human primate embryos^58^ (Fig. 2, Supplementary Note 1, and Supplementary Fig. 2 Dataset 1-11). In standard alignment benchmarks characterized by identical platforms and high tissue overlap (Fig. 2a), we subjected each dataset to five independent rounds of random spatial rotations and coordinate perturbations (Supplementary Fig. 3; see Methods). MAPS-alignment consistently achieved stable, accurate, and reproducible alignments across all biological specimens and perturbation trials within s (execution time of 10.59 ± 9.42 s for all rounds and all datasets), requiring a peak GPU memory footprint of merely ∼0.02 GB (Fig. 2a and Supplementary Fig. 3-10). Quantitative evaluations further demonstrated that MAPS-alignment achieved a highly favorable balance between alignment accuracy and resource efficiency while maintaining robust performance (Fig. 2b and Supplementary Note 9).

**Fig. 2.**
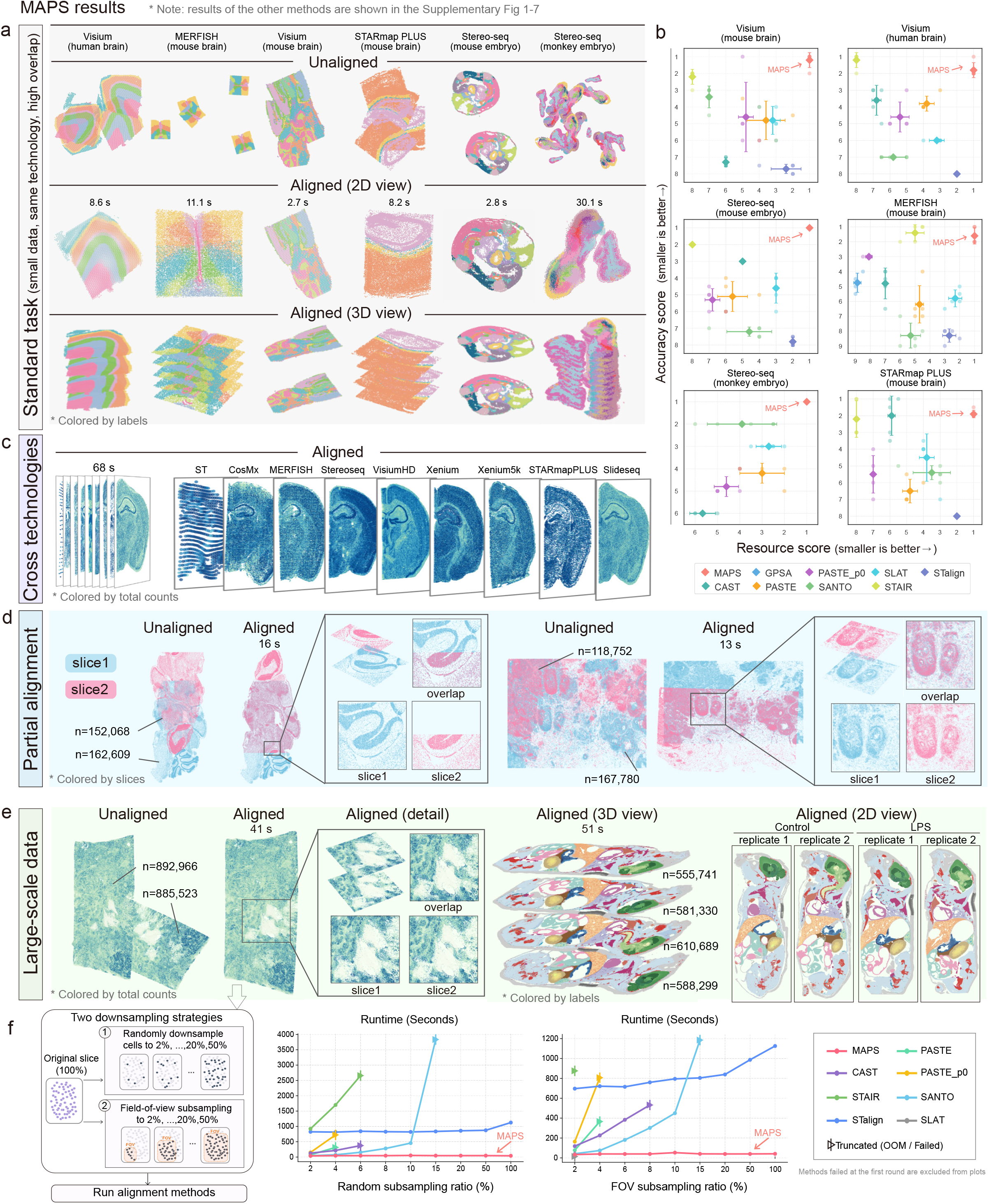
MAPS enables accurate and resource-efficient spatial alignment across diverse scenarios. **a**, Standard-scenario alignment using MAPS. Each column represents one dataset, with technology and tissue region indicated above. Raw slices and alignment results in 2D and 3D are shown from top to bottom. Each dataset was randomly shuffled and evaluated in five independent runs; additional results are shown in Supplementary Fig. 3–9. **b**, Quantitative evaluation of alignment accuracy and computational efficiency (five times independent run). The resource score integrates runtime, CPU memory, and GPU memory usage (Methods), with lower values indicating better performance. The y-axis represents the alignment accuracy score, again with lower values indicating better performance. **c**, MAPS alignment across sequencing techniques. Each slice in this dataset originates from a different platform. Compressed and expanded side views of the aligned volume are shown on the left and right sides, respectively. Slices are colored by total counts using independent scales. **d**, MAPS alignment under partial spatial overlap. Raw and aligned views are shown for each dataset, with magnified insets, spot numbers and runtimes indicated. **e**, MAPS alignment of large-scale spatial data, with spot numbers and runtimes indicated. **f**, Down sampling analysis of the dataset in **e** using random and FOV-based sampling. Runtime across sampling ratios is shown for each method. Additional results for more challenging scenarios are provided in Supplementary Figs. 11–14.

We next evaluated the robustness of MAPS-alignment under several challenging real-world analytical scenarios, including cross-platform alignment (Fig. 2c), partial slice overlap (Fig. 2d), and ultra-large-scale datasets (Fig. 2e). When aligning mouse brain coronal slices acquired across nine distinct spatial platforms (Supplementary Fig. 2 Dataset 7 and Supplementary Fig. 11a), MAPS-alignment uniquely achieved precise spatial alignment (Fig. 2c). In contrast, alternative algorithms struggled to produce reliable alignments, largely due to drastic discrepancies in spatial resolution, spot density, and gene panel coverage across platforms (Supplementary Fig. 12a, column 2). Furthermore, in partial alignment tasks where only a fraction of the tissue was shared between slices, for example because of uneven tissue trimming or fractured slices (Supplementary Fig. 2 Dataset 8-9), MAPS-alignment achieved accurate alignment of the overlapping regions (Fig. 2d), whereas other approaches either failed to complete the alignment (Supplementary Fig. 12a, column 3 and 4) or yielded substantial misalignments (Supplementary Fig. 12b, left).

Crucially, MAPS-alignment also exhibits outstanding computational scalability, effortlessly processing large-scale datasets containing millions of cells. For slice pairs with hundreds of thousands of cells, MAPS-alignment completed alignment on both Dataset 8 (Fig. 2d, left, Supplementary Fig. 2) and 9 (Fig. 2d, right, Supplementary Fig. 2) in tens of s (mean runtime = 14.5 s) with a peak GPU memory consumption of ∼3 GB (mean = 3.05 GB) (Fig. 2d and Supplementary Fig. 2 the last two columns), achieving a 104.86-fold speedup over STalign^51^ (the only other method that completed the task on both Datasets 8 and 9) (Supplementary Fig. 12b, columns 3 and 4), which required 25.34 min. For ultra-large-scale datasets (Fig. 2e; Supplementary Fig. 2, Datasets 10-11), MAPS-alignment completed alignment in under one minute (mean runtime = 46.16 s) while maintaining peak GPU memory under 5 GB (mean = 3.84 GB). In contrast, all other benchmark methods either exceeded the 48-hour runtime limit or encountered OOM errors on the same 81-GB GPU hardware under identical data loads (Supplementary Fig. 12a, columns 5 and 6), whereas runs that completed produced substantial misalignments (Supplementary Fig. 12b, right). To evaluate algorithmic performance across varying data sizes, we tested both random and field-of-view (FOV) down sampling schemes with cell retention rates ranging from 2% to 100% (Fig. 2f; Supplementary Note 5: Random sub sampling and Field-of-view down sampling). As the fraction of retained cells increased, MAPS-alignment maintained low execution times and stable alignment performance under both down sampling strategies (Supplementary Figs. 13, 14). By comparison, alternative methods either failed to complete alignment at some retention rates or produced inaccurate alignments (Supplementary Figs. 13, 14). Together, these benchmarks demonstrate that MAPS-alignment provides an accurate, ultra-fast, and highly scalable rigid alignment foundation for intra-modal spatial biology.

### MAPS-alignment enables cross-modal alignment without shared features

Reconstructing multi-layered spatial tissue architecture requires aligning orthogonal modalities (such as epigenomics, transcriptomics, translatomics, proteomics, and metabolomics) and histology within a unified physical coordinate system. However, existing computational alignment algorithms^42,50-55^ fundamentally rely on predefined molecular correspondences between modalities, for example links between accessible chromatin regions and expressed genes, or shared genes between transcriptomic and translatomics measurements, which limits their applicability when such correspondences are absent (e.g., between spatial epigenomics and metabolomics, or between transcriptomics and image data). Furthermore, specialized tools may require manual specification of rotation angles and may not automatically estimate coordinate scaling across platforms (e.g., SpatialMETA^59^). To overcome these limitations, MAPS-alignment utilizes a geometry-driven spatial alignment strategy that completely decouples physical coordinates from molecular feature dependencies, enabling fully automated alignment across arbitrary spatial platforms without requiring shared molecular features.

To evaluate the capacity of MAPS-alignment to align spatial data without shared molecular features, we first analyzed three colon adenocarcinoma (COAD) datasets^60^ (Supplementary Fig. 2 Dataset12-14) generated across distinct high-resolution spatial platforms (Xenium 5k, CosMx 6k, and Visium HD) each with a corresponding adjacent slice profiled by CODEX spatial proteomics (Fig. 3a). Given the multi-million cell scale and the absence of shared features between the cross-modal slice pairs, MAPS-alignment was the only tested method capable of processing this dataset. MAPS-alignment aligned all 6 slices across the four distinct platforms and modalities within a unified physical coordinate system in 150 s (Fig. 3b), while competing tools failed to execute. Beyond adjacent slices, MAPS-alignment also successfully resolved spatial alignments across diverse multi-slice combinations, including adjacent and non-adjacent cross-omics slices as well as non-adjacent intra-omics slices (Fig. 3c). We further tested the sensitivity of MAPS-alignment on a mouse dorsal tongue dataset^61^ characterized by subtle morphological boundaries (Fig. 3d and Supplementary Fig. 2 Dataset15). MAPS-alignment estimated coordinate scaling factors (scaling factor ≈ 1.958) and rotational angles (rotation ≈ 123°), achieving alignment of lymphoid and epithelial compartments (Fig. 3d).

**Fig. 3.**
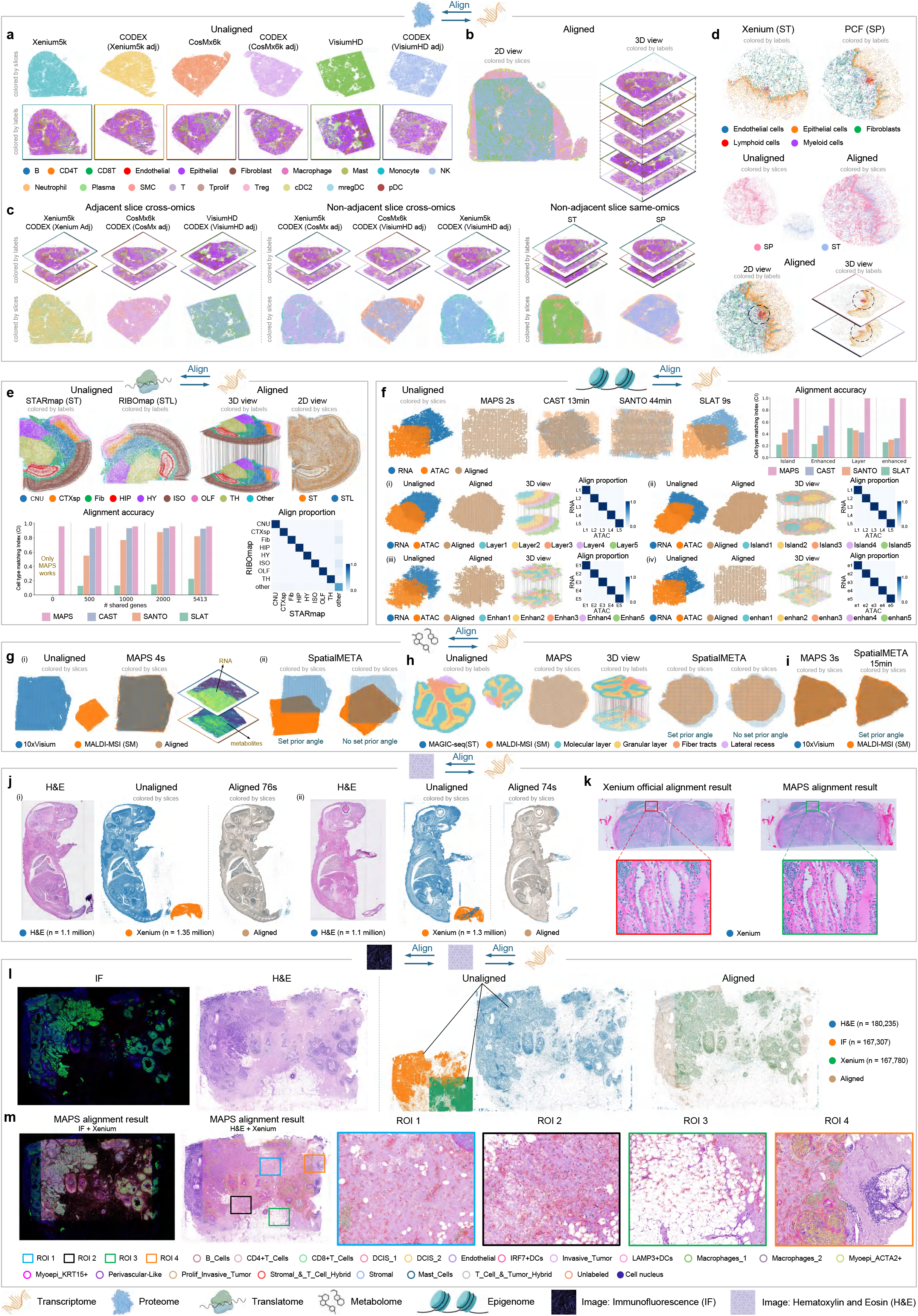
MAPS enables cross-modal alignment without shared features. **a**, Illustrations of colon adenocarcinoma slices profiled by three spatial transcriptomics platforms and their adjacent CODEX spatial proteomics slices. **b**, 2D (left) and 3D (right) visualizations of the six slices after MAPS alignment. **c**, MAPS alignment on cross-omics adjacent, cross-omics non-adjacent and same-omics non-adjacent scenarios. **d**, Cross-modal alignment of mouse dorsal tongue slices profiled by MERFISH and PCF. We illustrated spatial distributions before and after alignment and 3D visualization of aligned lymphoid cells. **e**, Alignment of coronal mouse brain slices. The alignment proportion matrix quantifies spatial nearest-neighbor consistency, and the bar plot compares cell-type matching accuracy across methods across different numbers of shared features. **f**, Benchmarking epigenomic–transcriptomic alignment using four distinct scMultiSim simulated datasets (I, ii, iii, iv). Alignment results and the corresponding nearest-neighbor consistency matrices are shown. **g**, Aligning spatial transcriptomics and metabolomics human liver cancer data using MAPS and SpatialMETA. (i) Original illustrations of the two slices, and the alignment results from MAPS with spatial visualization of transcriptomic (COL5A3) and metabolic (m/z 344.0562) features. (ii) Alignment results of SpatialMETA in two settings. **h**, Comparison of MAPS and SpatialMETA for mouse cerebellum alignment, with SpatialMETA evaluated with and without prior rotation information. **i**, 2D alignment results of human renal cell carcinoma from MAPS and spatialMETA. **j**, Alignment of hematoxylin and eosin (H&E) stained images with Xenium transcriptome dataset in mouse embryo; the H&E image was segmented into cell nuclei using Cellpose. **k**, Alignment of H&E images with Xenium transcriptome dataset in human renal cell carcinoma: comparison between MAPS and Xenium official results. **l**, The H&E image, immunofluorescence (IF) image and Xenium transcriptome dataset from human breast cancer before and after alignment. **m**, Overlay of aligned H&E, IF and Xenium data, with Xenium coordinates colored by cell type. Zoomed-in regions show that MAPS successfully aligned Xenium transcriptome dataset and H&E `with high accuracy`.

We next benchmarked MAPS-alignment directly against existing spatial alignment tools (SLAT^52^, CAST^40^, and SANTO^53^) in settings where the molecular correspondences required by these methods were available, including shared genes between transcriptomic and translatomics data and gene-level links between chromatin accessibility and gene expression. When aligning RIBOmap translatomics and STARmap transcriptomics coronal slices of the mouse brain^40^ using all 5,413 shared features (Supplementary Fig. 2 Dataset16), SLAT, SANTO, and CAST achieved relatively high Cell type matching index (CI) (Supplementary Note 9) scores of 0.23, 0.82, and 0.93, respectively, all below the 0.96 achieved by MAPS-alignment. As we progressively down sampled the shared features from 5,413 to zero, the performance of SANTO and SLAT declined substantially, with their CI scores decreasing by 30% and 50%, respectively, whereas MAPS-alignment maintained robust alignment performance even in the absence of shared features (Fig. 3e and Supplementary Fig. 15a). The nearest neighbor alignment ratio (Supplementary Note 9) of MAPS also exceeded 0.9 across all annotated anatomical regions except the category labelled “Other”.

To evaluate the alignment performance of MAPS on epigenomic and transcriptomic data, we used the multi-omics simulated dataset from SPCoral^62^, which comprises four spatial distribution patterns simulated by scMultiSim^63^ (Supplementary Fig. 2 Dataset17-20). Notably, these benchmark datasets were independently generated by the original authors of SPCoral rather than designed by us, ensuring an impartial and unbiased evaluation. MAPS-alignment achieved a 100% alignment success rate, accurately recovering ground-truth rotational angles (mean error = 0°) and translational displacements (mean error = 0 μm) across all four simulated datasets (Fig. 3f and Supplementary Fig. 15b). In contrast, CAST, SLAT, and SANTO performed poorly across all simulation runs. Furthermore, MAPS-alignment achieved successful alignment between spatial metabolomics and spatial transcriptomics in human liver cancer^64^, mouse cerebellum^25^, and human renal cell carcinoma (RCC)^65^ (Fig. 3g-I, Supplementary Fig. 2 Dataset21-23). While SpatialMETA^59^ represents a specialized metabolome alignment tool, it requires manual specification of rotation angles and the learned coordinate scaling factors perform poorly, resulting in residual positional misalignments (Fig. 3g, h and Supplementary Fig. 15c-e). To evaluate the effect of rotation-angle initialization in SpatialMETA, we tested it with and without prior rotation angle specification. In contrast, MAPS-alignment automatically resolved scaling and rotational offsets, achieving a 120-fold speedup (3 s vs. 15 min, Fig. 3i). Crucially, MAPS-alignment achieved a CI of 0.75 without a prior rotation angle, surpassing both SpatialMETA configurations: 0.36 (no prior angle) and 0.7 (prior angle) (Supplementary Fig. 15e).

Finally, we evaluated MAPS-alignment on large-scale tasks involving image alignment. In aligning two pairs of whole-embryo H&E images against single-cell resolution Xenium transcriptomic datasets containing over one million cells (Supplementary Fig. 2 Dataset24, 25), MAPS-alignment completed the alignment in approximately 74 s with a peak GPU memory consumption of 4.9 GB (Fig. 3j). To evaluate alignment fidelity against a manual proofreading, we aligned human renal cell carcinoma slices (Supplementary Fig. 2 Dataset26) and compared regions of interest (ROIs) directly against alignments generated by Xenium official, which requires extensive manual landmark pinning (Fig. 3k). The spatial coordinates aligned by MAPS showed visual agreement with the manually proofread results (Fig. 3k). We also applied MAPS-alignment to tri-modal spatial data from human breast cancer^9^ (Supplementary Fig. 2 Dataset27), aligning H&E histology, IF, and Xenium in situ RNA profiling within the same tissue block (Fig. 3l). High-resolution magnification views of four representative ROIs demonstrated that the aligned Xenium transcript coordinates mapped directly to corresponding cell nuclei identified by DAPI and H&E staining (Fig. 3m). Together, these results demonstrate that MAPS-alignment provides a fully automated, scalable, and accurate solution for cross-modal alignment without requiring shared molecular features or manual intervention.

### MAPS enables cross-modal integration across anchored and unanchored scenarios

Integrating spatial information across multiple modalities, including transcriptomics, proteomics, metabolomics, and histology, is essential for characterizing complex tissue biology. Built upon physically aligned coordinate spaces, MAPS further introduces two specialized integration engines: MAPS-integration-U enables anchor-free diagonal integration by propagating molecular information across spatially aligned tissue sections; MAPS-integration-P uses available histological images as structural anchors to transfer multi-modal information across tissue volume.

To validate MAPS-integration-U, we first applied it to consecutive colon adenocarcinoma (COAD) slices profiled by spatial transcriptomics (Xenium 5k, CosMx 6k, and Visium HD) and spatial proteomics (CODEX) (Fig. 4a, Supplementary Fig. 2 Dataset12-14). The results demonstrated that MAPS-integration-U effectively integrates two arbitrary modalities, revealing consistent spatial domain patterns across both adjacent (Fig. 4b, (I, ii)) and non-adjacent section combinations (Fig. 4b, (iii)). We further applied MAPS-integration-U to adjacent spatial transcriptomic and proteomic sections of the mouse dorsal tongue (Supplementary Fig. 2 Dataset15), resolving the manually annotated epithelial-enriched domain into two functionally distinct sub-domains 0 and 2 (Fig. 4c-e). Domain 0 was characterized by basal/progenitor markers (*Tp63*^66^, *Krt6a*^67^ and *Egfr*^68^), high proliferative protein activity Ki67^69,70^ and Histone H3 Phospho (Ser28)^71^, suggesting its role as a progenitor-rich proliferative compartment (Supplementary Fig. 16a-c). In contrast, domain 2 exhibited differentiated epithelial features *Krt4*^67,72^, enriched immune recruitment signals *Mal*^*73*^, expression of adhesion and immunomodulatory protein CD66a^74^, and a balanced inflammatory state (*Slpi*^75^ and *Il1rn*^76^), indicating a role in immune surveillance at the epithelial interface (Supplementary Fig. 16d-f). Together, these results demonstrate that MAPS-integration-U resolves spatially regionalized functional subdomains by integrating unanchored orthogonal modalities.

**Fig. 4.**
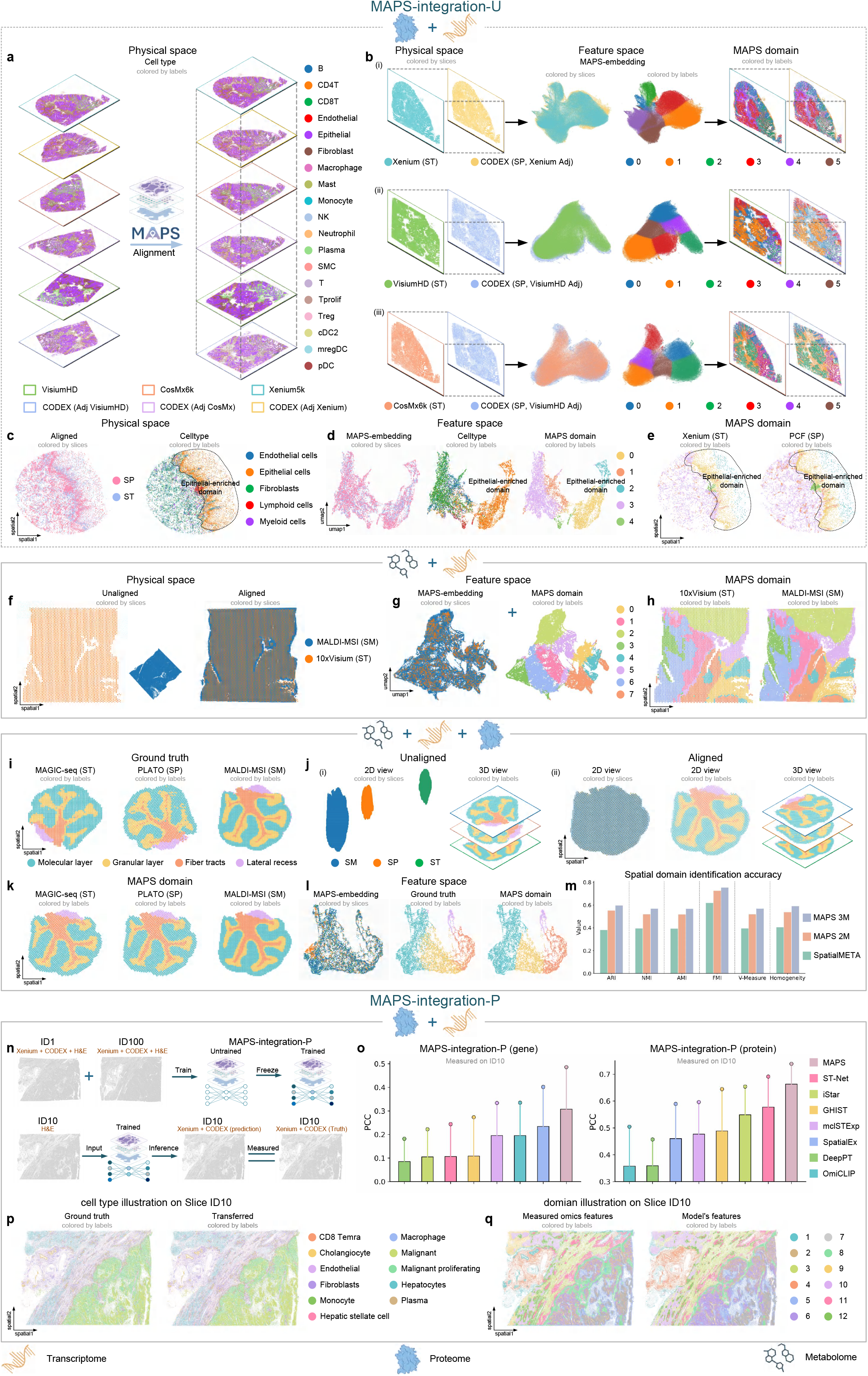
MAPS enables unanchored diagonal integration and anchored 3D multi-modal integration. **a**, MAPS-alignment achieves multi-slice alignment of `unanchored` colon adenocarcinoma. **b**, (i-iii) Alignment `result` of unanchored cross-modal slices (left). UMAP visualization of the integrated transcriptome and proteome datasets (middle). Each cell is colored by omics or spatial domain identification. Spatial mapping of domain identification results (right). **c**, Alignment results of the mouse dorsal tongue. **d**, UMAP visualization of the integrated transcriptome and proteome in the mouse dorsal tongue. Each cell is colored by omics types (left), cell type annotations (center) and identified spatial domains (right). **e**, Spatial mapping of spatial domain identification results. **f**, Visualization of MAPS alignment results in human liver cancer. **g**, UMAP visualization of the integrated transcriptome and metabolome. Each cell is colored by different platforms (left) and domain identification result (right). **h**, Spatial mapping of spatial domain identification results. **i**, Spatial mapping of the mouse cerebellum before alignment. Each spot is colored by manual annotation. **j**, (i) 2D and 3D view of the mouse cerebellum before alignment. Each spot is colored by modal (left) and manual annotation (right). (ii) 2D and 3D view of the mouse cerebellum after alignment. Each spot is colored by modal (left) and manual annotation (center and right). **k**, Spatial mapping of the mouse cerebellum after alignment. Each spot is colored by spatial domain identification results. **l**, UMAP visualization of the integrated transcriptome, proteome and metabolome. Each cell is colored by different omics (left), manual annotation (center) and spatial domain identification result (right). **m**, Barplot of quantitative evaluation for 3 modal MAPS-integration-U diagonal integration, 2modal MAPS-integration-U diagonal integration and SpatialMETA with 6 supervised metrics. **n**, MAPS-integration-P was trained using anchored multi-omics data of ID1 and ID100 and evaluated using ID10. **o**, Omics prediction performance of MPAS-integration-U. **p**, Cell type transfer results based on measured omics features and cross-slice transfer from slices ID1 and ID100 to ID10. **q**, Spatial domain identification using measured omics data compared with results from the model’s features.

We next aligned and integrated two adjacent liver cancer sections^64^ profiled by MALDI-MSI and 10x Visium, respectively (Fig. 4f-h and Supplementary Fig. 2 Dataset23). The spatial patterns of differentially expressed metabolic and transcriptional markers were concordant with the identified domains (Supplementary Fig. 16g), demonstrating that MAPS-integration-U can integrate adjacent multimodal data across distinct spatial resolutions while preserving spatially coherent tissue organization. To further evaluate its diagonal integration capability, we extended MAPS-integration-U to triple-modal integration and benchmarked it against SpatialMETA^59^ using mouse cerebellum data^25^ (Supplementary Fig. 2 Dataset22). MAGIC-seq, PLATO, and MALDI-MSI profiled sections were spatially aligned and jointly integrated (Fig. 4i, j), and the resulting latent representations were used for spatial domain identification (Fig. 4k, l). For direct comparison with SpatialMETA, we additionally performed bimodal integration using MAGIC-seq and MALDI-MSI data (Fig. 4m). The results demonstrated that MAPS with triple-modal integration achieved the highest scores across six evaluation metrics. MAPS-integration-U with bi-modal integration ranked second, outperforming SpatialMETA by more than 20% (Adjusted Rand Index (ARI) of 0.59 for triple-modal MAPS-integration-U, 0.55 for bi-modal MAPS-integration-U, and 0.38 for SpatialMETA).

To assess MAPS-integration-P for histology-anchored multi-modal integration, we conducted the experiments using our in-house human hepatocellular carcinoma (HCC) dataset comprising serial tissue sections ordered along the z-axis (ID1 to ID100; see Methods). Among these, three sections positioned at different depths (ID1, ID10, and ID100) were jointly profiled by single-cell-resolution spatial transcriptomics, spatial proteomics, and H&E imaging, while the remaining sections contained H&E staining alone. We trained the histology-based multi-omics prediction model on sections ID1 and ID100, and evaluated its generalization on the intermediate held-out section ID10 (Fig. 4n). Quantitative evaluation demonstrated that MAPS-integration-P substantially outperformed existing state-of-the-art methods^49,77-82^ in predicting gene expression and protein expression from H&E images at cellular resolution (Fig. 4o). Compared with the second-best methods (SpatialEx for gene expression prediction and ST-Net for protein abundance prediction), MAPS-integration-P improved the mean performance by 31% and 14%, respectively. Furthermore, MAPS-integration-P supported downstream biological characterization on the held-out ID10 section, and the predicted cell types (Fig. 4p) and inferred spatial domains (Fig. 4q) closely recapitulated their corresponding reference standards (see Methods).

### MAPS-Explorer provides interactive 3D analysis and visualization capabilities for spatial biology

At whole-organ or organism scale, interactive visualization of datasets comprising millions to tens of millions of cells remains computationally challenging. Although Spateo provides the browser-based Spateo-viewer^83^, its scalability to tens of millions of cells has not been systematically characterized, and the extent to which native 3D interaction can facilitate biological discovery beyond conventional 2D analysis remains unclear. We therefore developed MAPS-Explorer, an open-source, self-hosted platform that supports cross-system (Windows, macOS, Linux), cross-device, and multi-modal visualization, along with diverse native spatial interactive analysis functions for large-scale spatial omics data (Fig. 5a, b and Supplementary Note 2). MAPS-Explorer provides a user-friendly interactive interface together with a customized rendering pipeline, reducing both the lower the barrier to use and hardware requirements. To further facilitate adoption, we provide a user manual of more than 30 pages, step-by-step video tutorials, multiple example datasets, and a publicly accessible interactive website https://bioinfor.imu.edu.cn/maps-explorer/.

**Fig. 5.**
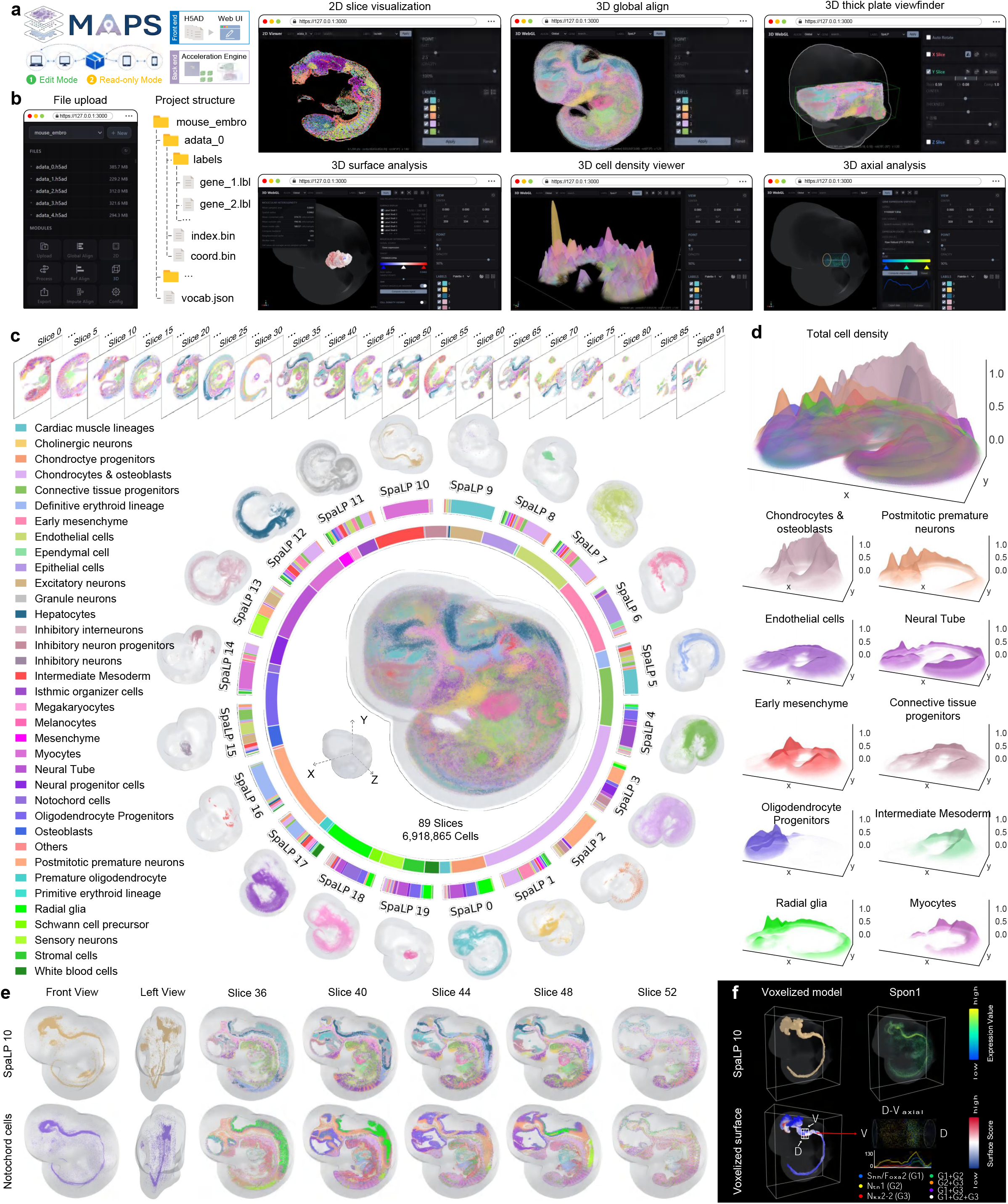
MAPS-Explorer, an open-source platform for interactive 3D visualization of spatial omics data. **a**. MAPS-Explorer supports cross-platform deployment across major operating systems and devices. **b**. MAPS-Explorer provides a comprehensive suite of visualization and interaction modules, including File upload, 2D slice visualization, 3D global alignment, 3D thick plate viewfinder, 3D surface analysis, 3D cell density viewer and 3D axial analysis, together with highly customizable visualization and filtering parameters. **c**. Reconstruction of an E11.5 mouse embryo from 89 consecutive spatial transcriptomic slices obtained from Spateo. The outer and inner rings summarize cell-type composition from SpaLP-defined spatial domains and across the entire embryo, respectively. The central whole-embryo and surrounding domain-resolved 3D models are colored by SpaLP annotations. **d**. 3D cell-density visualization of the whole embryo and the ten most abundant cell types. **e**. 3D reconstruction reveals the continuous notochord structure from both SpaLP and cell-type distributions, which is difficult to appreciate in individual 2D slices. **f**. The notochord structure was used to build a voxelized model and a voxelized surface. Both Spon1 expression and surface score exhibit a strong co-localization of the notochord. A vertical cylindrical space passing through the dorsal notochord region is used to screen the axial cells. We rotated the cylinder counterclockwise by 90 degrees as the observation direction. The left side represents the dorsal side, and the right side represents the ventral side. Furthermore, we quantified the co-expression of the dorsal-related gene Nkx2-2 (shown in the red line graph), the notochord marker gene Ntn1 (shown in the yellow line graph) and the ventral-related gene Shh/Foxa2 (shown in the blue line graph).

MAPS-Explorer leverages a Python-based core and a decoupled frontend–backend architecture to combine native 3D surface, axial, colocalization, and cell-interaction analyses with a proprietary acceleration engine and WebGL-based rendering. It enables real-time interactive querying and rendering of datasets containing tens of millions of cells (Fig. 5a, b and Supplementary Fig. 17a; see Methods). To quantitatively evaluate the interactive performance of MAPS-Explorer, we benchmarked loading, and rendering across datasets and deployment configurations (Supplementary Fig. 17b). MAPS-Explorer remained stable across both local and remote deployments. Loading time increased approximately linearly as dataset size increased from ∼0.8 million to >30 million cells, whereas interactive frame rates remained largely independent of dataset size. Moreover, MAPS-Explorer supports real-time filtering and rendering of arbitrary cell labels across multiple spatial levels, spanning whole-tissue organization, spatial domains, and individual cell types, while preserving continuous spatial structures across four datasets (Supplementary Fig. 17c–e). For the 4 benchmark datasets (Supplementary Fig. 2 Dataset 29-32), spatial continuity after alignment was assessed using the original annotation. Consequently, the continuous structures observed along the z-axis directly reflect the preservation of pre-existing anatomical and cellular organization after alignment (Supplementary Fig. 17e and Supplementary Note 10).

We applied MAPS to reconstruct an E11.5 mouse embryo^83^ from 89 serial spatial transcriptomic slices and explored the 3D model in MAPS-Explorer. The atlas comprised 6,918,865 cells, 21,717 genes, 20 continuous spatial domains identified by SpaLP^84^ and 36 major tissues or cell types (Fig. 5c; see Methods, Supplementary Fig. 2 Dataset29), and could be loaded and rendered in approximately 5 s (deployed locally). An enhanced Marching Cubes^85^ construction pipeline was used to generate continuous 3D voxelized shells from point clouds (see Methods). The concentric annotation rings summarized cell-type composition at the domains and whole-embryo scales. The domain-resolved models separately display the 3D morphology and anatomical location of each spatial domain (Fig. 5c). MAPS-Explorer revealed smooth and spatially continuous alignment across both the spatial-domain and cell-type levels. The outer ring shows the relative cell-type composition within individual spatial domains, whereas the inner ring summarizes cell-type composition across the entire embryo. The surrounding domain-resolved models enable independent visualization of the 3D morphology, spatial extent, and anatomical location of each spatial domain (Fig. 5c). The dense distribution of Chondrocytes & osteoblasts and Postmitotic premature neurons along the dorsal side reflected active development of vertebral and neural structures at E11.5^86^ (Fig. 5d).

Notably, spatial domain 10 is a narrow continuous domain along the embryonic midline corresponding to the notochord. Because of its slender morphology and the oblique orientation of the embryo during sectioning, the notochord appeared as spatially disconnected cross-slices in individual 2D slices, obscuring its longitudinal continuity. By contrast, 3D visualization clearly recovered the complete axial structure (Fig. 5e and Supplementary Fig. 18a, b). Within the notochord region, midbrain neuroectoderm, cranial mesoderm and epithelial precursors predominated, while *Shh*^58^, *En2*^87^ and *Nkx6*^88^ showed spatially restricted expression patterns consistent with axial patterning and notochord-related developmental programs (Supplementary Fig. 18a–d). Finally, we reconstructed the notochord surface and quantified the spatial heterogeneity of *Spon1*^89^ along its dorsoventral axis, revealing a pronounced axial asymmetry (Fig. 5f). 3D axial analysis combined with 3D colocalization resolved two spatially adjacent but distinct molecular domains, comprising a *Shh*^+^/*Foxa2*^+90^ axial midline domain and a dorsally displaced *Nkx2-2*^+91^ neural domain. Both domains were associated with *Ntn1* expression but formed discrete peaks along the dorsoventral axis.

Taken together, MAPS-Explorer enables efficient and interactive analysis of spatial datasets comprising tens of millions of cells and facilitates the visualization of continuous 3D anatomical structures that are difficult to resolve from individual 2D slices. It further enables quantitative characterization of spatial organization and developmental patterning through integrated surface, axial and colocalization analyses.

### MAPS constructs a multi-modal 3D mouse brain atlas

The mammalian brain possesses a highly stereotyped anatomical architecture that has long provided a stable spatial scaffold for cross-specimen data integration, as demonstrated by classical stereotaxic coordinate frameworks and connectome atlases in which data from individual brains are registered onto a common coordinate system^92,93^. While international consortia such as the BRAIN Initiative Cell Census Network aim to systematically map the spatial cellular landscape of the brain, no single spatial profiling technology can simultaneously capture all molecular modalities at the whole-brain scale within a single specimen. Existing brain spatial atlases^6,94-98^ therefore remain fragmented as isolated datasets defined by distinct probe panels^16^, disparate spatial resolutions, and incompatible local coordinate systems, preventing integrated cross-modal analysis. To resolve these limitations and establish a standardized spatial consensus, we applied MAPS to assemble the first unified, multi-modal 3D spatial coordinate framework for the mouse brain, bridging 18 independent atlases comprising 5 modalities, 434 coronal sections, and 21,335,013 cells / spots (Fig. 6a), including Chen et al.^94^ (Atlas 1, BARseq), Shi et al.^96^ (Atlas 2, STARmap PLUS), Zhang et al.^98^ (Atlas 3, MERFISH), Yao et al.^97^ (Atlas 4, MERSCOPE), Vizgen. (Atlas 5, MERSCOPE), 10xGenomics. (Atlas 6, Xenium), 10xGenomics. (Atlas 7, Xenium5k), 10xGenomics. (Atlas 8, Xenium), STOmics. (Atlas 9, Stereo-seq), NanoString. (Atlas 10, CosMx), Vizgen. (Atlas 11, MERFISH V2), Zeng et al.^19^ (Atlas 12, STARmap), Zeng et al.^19^ (Atlas 13, RIBOmap), Han et al.^95^ (Atlas 14, Stereo-seq), Han et al.^95^ (Atlas 15, Stereo-seq), Zhang et al.^30^ (Atlas16, CODEX), METASPACE. (Atlas17, MALDI-MSI), 10xGenomics. (Atlas18, H&E).

**Fig. 6.**
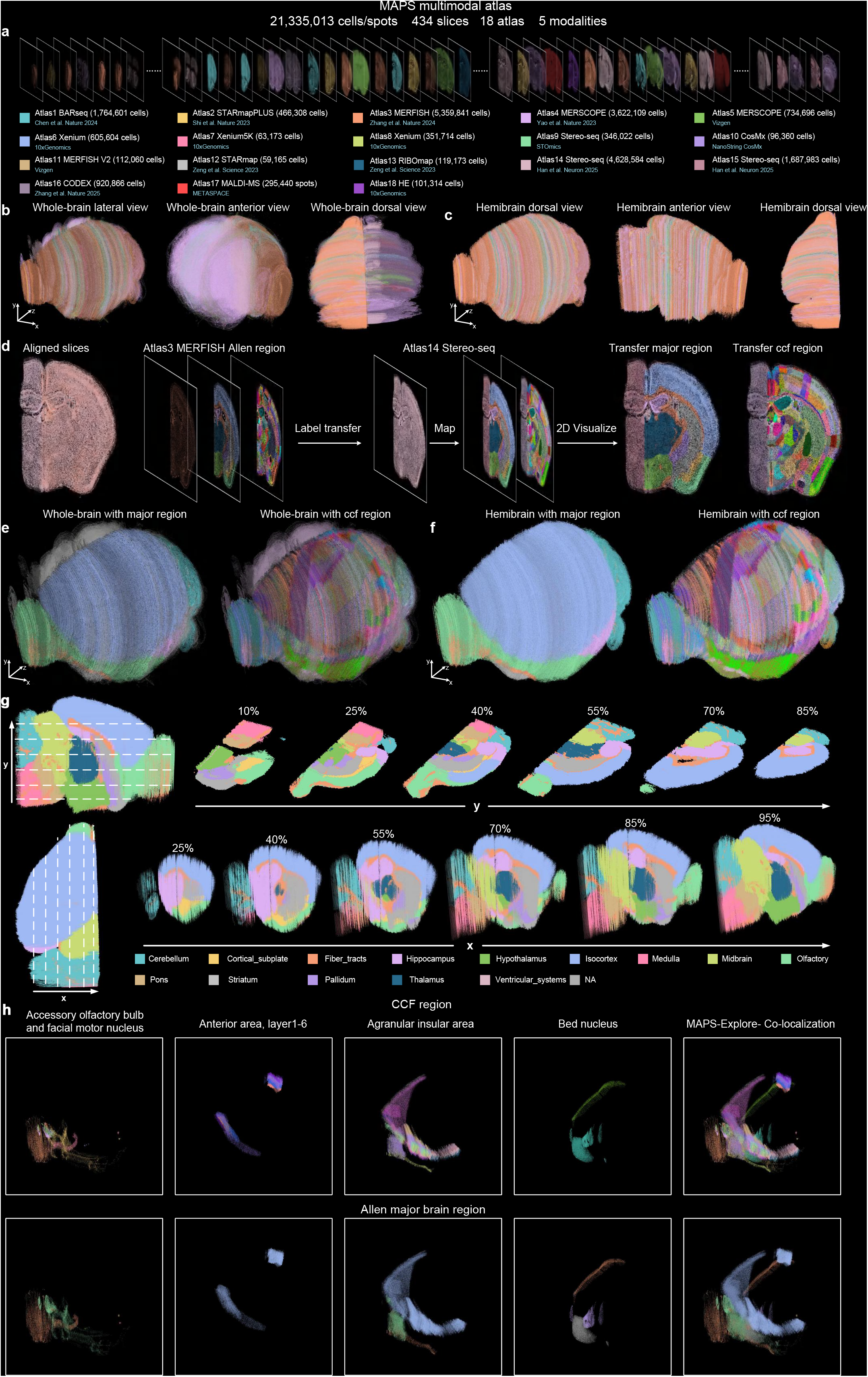
MAPS constructs the most comprehensive spatial multi-modal 3D atlas of the mouse brain. **a**, 3D spatial mapping of the coronal mouse brain slices across 5 modalities from 18 atlases. **b,c**, Spatial multi-modal 3D whole-brain (b) and hemi-brain (c) anatomical maps reconstructed by MAPS from 434 aligned slices, with cells/spots colored by atlas. **d**, MAPS enables cross-modal label transfer. Slices from other modalities were aligned to the MERFISH atlas, and Allen Brain annotations were transferred based on nearest-neighbor labels. **e**, Spatial multi-modal 3D whole brain anatomy perspective. Each cell/spot is colored by Allen major region (left) and Allen ccf region (right). **f**, Spatial multi-modal 3D hemibrain anatomy perspective. Each cell/spot is colored by Allen major region (left) and Allen ccf region (right). **g**, *In silico* sectioning using MAPS-Explorer. MAPS-Explorer was used to generate continuous in silico thick slices along the x-and y-axes. Each cell/spot is colored by Allen major region. **h**, Use the MAPS-Explorer hierarchical index to map the same cellular spatial information for the Allen major region and the ccf region, and visualize the co-localization of different hierarchical index spatial.

To integrate these diverse modalities into a unified spatial reference, we anchored the global coordinate system to the Allen Mouse Brain Common Coordinate Framework (CCF) defined by the MERFISH atlas in Atlas 3, aligning all coronal slices from the remaining 17 atlases to their corresponding stereotaxic positions (Fig. 6b, c). By decoupling physical coordinate alignment from molecular gene panel overlap, MAPS successfully aligned heterogeneous slices across vast differences in spatial resolution, from 0.2 μm cellular resolution to 50 μm tissue-level grids. Unlike conventional slice-by-slice alignment tools, which universally failed on these datasets, MAPS accomplished the global alignment of all 434 slices in under 1 hour with less than 5 GB of GPU memory. Leveraging this unified physical coordinate system, we developed a cross-modal label transfer pipeline to automatically propagate hierarchical Allen CCF anatomical annotations to the entire atlas (Fig. 6d). For any input slice, MAPS identifies spatial nearest neighbors within the aligned reference coordinate space, transferring annotations across two hierarchical tiers comprising coarse major brain regions and fine anatomical subregions. The inferred anatomical annotations exhibited high concordance with well-established histological annotations (Fig. 6d, right). Notably, this automated pipeline accurately delineated fine subcortical boundaries and preserved the spatial continuity of laminar structures across all modalities. This framework enabled the systematic, automated annotation of modalities that traditionally lack direct transcriptomic markers or comprehensive gene panels, including MALDI-MSI metabolomic profiles and low-plex CODEX proteomic slices, yielding a fully annotated, multi-modal 3D consensus atlas of the mouse brain (Fig. 6e–g).

While reconstructing a 3D organ volume provides holistic anatomical context^98^., understanding complex regional connectivity and boundaries requires flexible multi-angle visualization and internal tissue dissection. Current spatial biology workflows are typically constrained to fixed standard slicing planes (coronal or sagittal)^99^, which obscure oblique neuronal tracts and curvilinear structures. Although experimental sectioning along non-standard angles is physically possible, it requires extensive tissue sacrifice, increases experimental costs, and introduces confounding inter-individual anatomical variations. To overcome this limitation, MAPS-Explorer incorporates a continuous 3D simulated re-sectioning engine based on an arbitrary-angle sliding thick plate function (Fig. 6g). Unlike existing tools that only support 2D planar slicing along fixed Cartesian axes^99^, MAPS-Explorer enables users to define virtual sectioning plates of customizable thickness and smoothly slide them along arbitrary spatial trajectories and oblique angles. Furthermore, single-structure rendering and hierarchical label indexing allow users to computationally isolate discrete brain structures, such as the hippocampal formation or deep cerebellar nuclei, and analyze cross-modal molecular gradients, metabolic colocalization, and cellular organization within non-planar anatomical spaces (Fig. 6h).

### Multi-modal 3D hepatocellular carcinoma reconstruction

While the preceding assembly demonstrates the capacity of MAPS to integrate multi-modal atlases at the cross-consortium level, extending continuous multi-omics profiling to individual laboratory-scale 3D specimens represents a distinct and formidable challenge. Profiling continuous multi-layered molecular landscapes across deep 3D tissue volumes remains economically prohibitive and technically demanding^100^, as volumetric imaging methods suffer from limited tissue penetration depth and restricted sequencing panels^18^, whereas exhaustive serial sectioning incurs unsustainable reagent costs^101^. Consequently, reconstructing continuous single-cell 3D spatial multi-modal across multiple molecular layers remains largely unachievable for standard laboratory workflows. By utilizing routine, low-cost histological imaging as a structural scaffold, MAPS-integration-P leverages morphological features to propagate rich multi-modal measurements from a small set of sequenced anchor slices to the entire tissue volume.

To demonstrate the efficacy of this framework on laboratory-scale 3D specimens, we applied MAPS-integration-P to an in-house human hepatocellular carcinoma (HCC) dataset spanning dense serial slices (designated ID1 to ID100) (Fig. 7a, Supplementary Note 3 and Supplementary Figs. 2 Dataset 34). In this experimental design, only two terminal slices (ID1 and ID100) served as multi-modal anchors profiled with cellular-resolution spatial transcriptomics and proteomics, whereas the remaining intermediate slices were profiled solely with standard H&E imaging. Harnessing the algorithmic modules benchmarked in preceding sections, the computational pipeline proceeded through four steps: (i) MAPS-alignment to establish a continuous physical coordinate framework across all slices and all modalities; morphological representation learning using a pathology foundation model to extract deep visual embeddings from segmented single cells; (iii) multi-modal inference via MAPS-integration-P to predict continuous transcriptional and protein expression across all intermediate slices; and (iv) joint embedding-based label propagation to transfer cell-type and spatial-domain identities throughout the entire tissue volume (Fig. 7a).

**Fig. 7.**
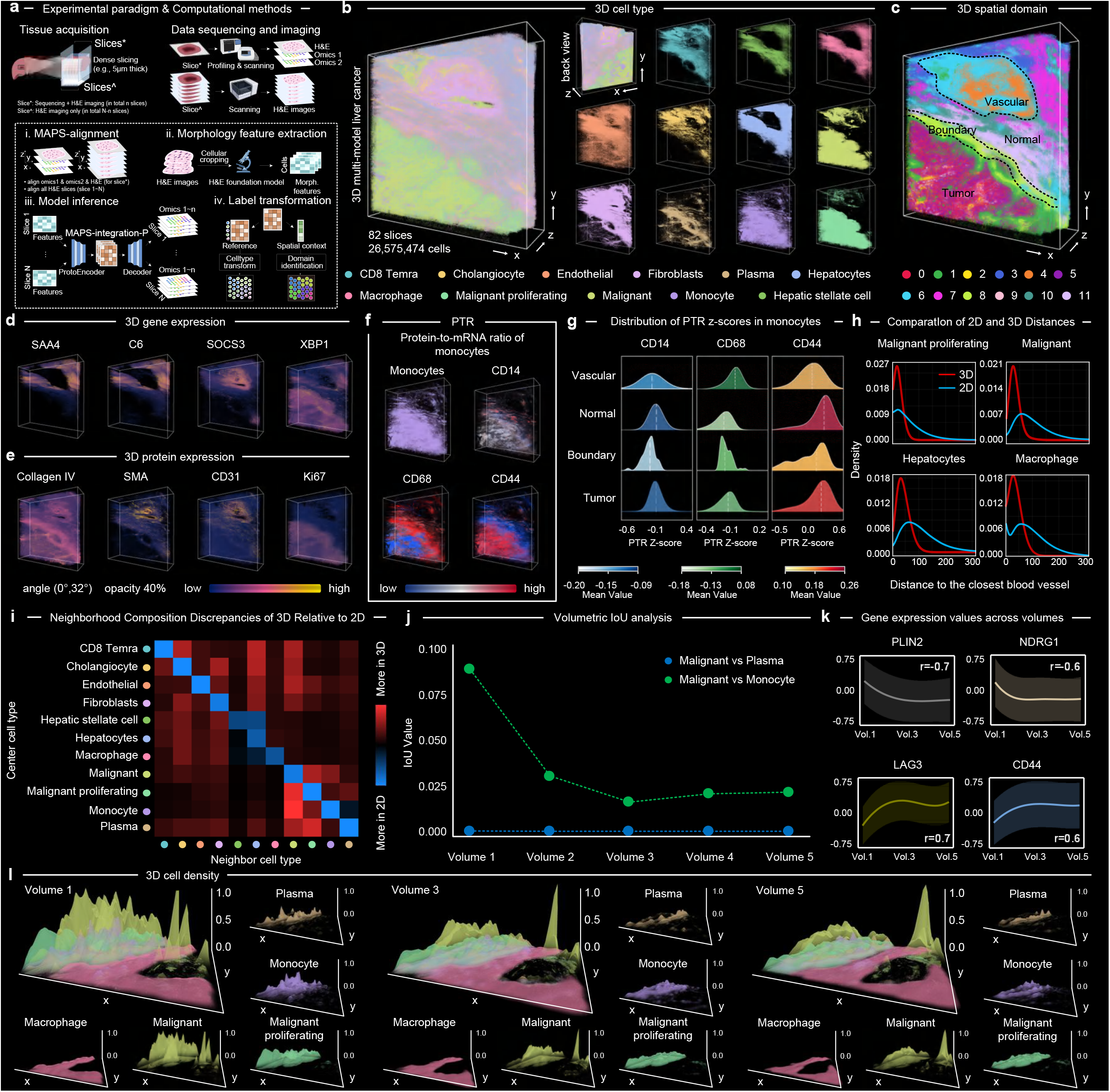
In-house Multi-modal 3D hepatocellular carcinoma reconstruction. **a**, Overview of the experimental paradigm (top) and MAPS-integration-P computation process (bottom). **b**, 3D visualization of the tissue volume in terms of mixed (right) and individual cell types (right). **c**, 3D spatial domain visualization and pathology annotation. **d-e**, 3D distributions of selected gene (d) and protein (e) signatures. **f**, Spatial distributions of monocytes and three protein-to-transcript ratio (PTR) scores. **g**, PTR scores of monocytes for CD14, CD68, and CD44 across tumor, boundary, normal, and vascular regions. Density curves indicate the median (white) and mean (colored). **h**, Density distributions of nearest-vessel distances are shown for the indicated cell types. Distances were computed in 2D within individual slices and in 3D using the full reconstructed vascular mask. Red and blue represent 3D and 2D distances, respectively. **i**, Differences in cell-type composition between 3D and 2D neighborhoods within a 20-μm radius. Rows and columns denote the centre and neighboring cell types, respectively; colors indicate differences between 3D and 2D proportions. **j**, Volumetric IoU values between malignant cells and plasma cells or monocytes. **k**, Expression values of monocyte-status-related marker genes across volumes. **l**, Gaussian-smoothed density maps and spatial overlap of important cell populations across volumes.

Following the previous validation (Fig. 4n-q), we reconstructed the 3D architecture of the tissue, incorporating multi-omics profiles, cell-type (Fig. 7b), and spatial domains (Fig. 7c and Supplementary Fig. 19a) (highly similar to pathological annotations outlined in black). We also examined the gene and protein markers and confirmed their consistency with the regional annotations (Fig. 7d, e and Supplementary Fig. 19b-d). Specifically, the proliferation marker Ki67 was highly expressed within the tumor region, while Collagen IV, a basement membrane component, was localized to the boundary. Vascular regions were characterized by high expression of SMA (a vascular smooth muscle cell marker^102^) and CD31(a vascular endothelial marker^103^), and inflammation-associated signatures (*SAA4*^104^, *C6*^105^, and *SOCS3*^106^) were enriched in the peritumoral area, suggesting a localized inflammatory state.

Leveraging the reconstructed 3D multi-omics volume, we investigated spatial variations in post-transcriptional regulation by quantifying the protein-to-mRNA ratio (PTR)^54^, which serves as an in-situ proxy for translational efficiency (Fig. 7f; see Methods). Given the dense spatial colocalization observed between infiltrating monocytes and malignant cells (Fig. 7b), we examined the spatial dynamics of PTR within the monocyte population across distinct microenvironments (Fig. 7f, g). Translational efficiency exhibited pronounced spatial heterogeneity across tissue compartments. The canonical monocyte marker CD14 showed a reduced PTR score within the tumor invasive boundary compared to adjacent non-tumor regions. Conversely, CD44 exhibited elevated PTR scores in both normal and malignant compartments, whereas CD68 PTR scores were significantly suppressed within the tumor core (Fig. 7g). These localized disparities in PTR uncover region-specific post-transcriptional reprogramming of monocytes within the tumor microenvironment that would remain entirely masked in single-modality transcriptomic analyses.

To investigate whether conventional 2D section analyses introduce systematic spatial distortions, we evaluated cellular geometric metrics across dimensions. We calculated the spatial distance from individual cell types to the nearest blood vessel in both 2D and 3D coordinate spaces (Fig. 7h, Supplementary Note 4, and Supplementary Fig. 19e). Across all cell types, 3D Euclidean distances to vasculature were consistently shorter than their 2D in-plane counterparts, with an average distance reduction of 50.27 ± 57.88 μm (P < 2.2 × 10^−16^, paired two-sided Student’s t-test), directly attributable to out-of-plane vessels traversing above or below individual 2D slices (Fig. 7h). Furthermore, cellular neighborhood analysis revealed that 2D projections significantly overestimated homotypic cell-cell clustering by 33.98 ± 21.09% (P = 8.22× 10^−5^, paired two-sided Student’s t-test across cell types), whereas 3D volumetric neighborhoods captured a substantially greater diversity of heterotypic cell-cell contacts (Fig. 7i). These findings demonstrate that planar 2D analyses introduce significant spatial truncation bias, underscoring the critical necessity of true 3D volumetric reconstruction for accurately characterizing cellular microenvironments.

Finally, we interrogated continuous volumetric cellular transitions along the z-axis by constructing Gaussian-smoothed density volumes for major cell populations across successive 20-slice sub-volumes (from volume 1 to volume 5, advancing deeper into the tumor mass) (Fig. 7j–l and Supplementary Fig. 19f). While intervening plasma cells physically segregated malignant cells from mature macrophages, monocytes exhibited dense colocalization with malignant cells in superficial layers (Fig. 7l). Volumetric Intersection over Union (IoU) analysis revealed a progressive attenuation of monocyte-malignant colocalization along the z-axis, declining monotonically from 0.817 in sub-volume 1 to 0.198 in sub-volume 5 (Fig. 7j). This spatial decoupling corresponded to a continuous z-axial transition from acute inflammatory signaling toward mature macrophage phenotypes in deeper tumor regions. This phenotypic evolution was further corroborated by marker dynamics: metabolic and hypoxic stress markers (*PLIN2* and *NDRG1*) were progressively downregulated along the z-axis, accompanied by the concomitant upregulation of immune checkpoint and adhesion regulators, including *LAG3* and *CD44* (Fig. 7k, l). Together, these spatial and molecular transitions delineate a directional progression from acute stress adaptation toward an immunosuppressive, tumor-associated macrophage (TAM) phenotype^107^ within deep tumor region.

## Discussion

The structural and functional principles of multicellular life are governed by the intricate interplay between a cell’s high-dimensional molecular state and its precise physical coordinates within native tissue architecture. While spatially resolved profiling technologies have revolutionized cellular cartography, the field has hitherto remained constrained by a flatland paradigm, fragmented across isolated 2D slices, disparate molecular modalities, and incompatible coordinate systems. MAPS provides the missing computational infrastructure to transcend these limitations, establishing a unified physical and molecular coordinate framework across arbitrary spatial platforms. Rather than assembling modality-specific solutions piecemeal, MAPS provides the missing computational infrastructure to transcend the flatland limitations of fragmented spatial data, establishing a unified physical and molecular coordinate framework across arbitrary platforms through 3 core modules that address the long-standing disconnect between physical registration and feature integration.

In this study, MAPS-alignment decoupled coordinate registration from shared molecular features, operating purely on tissue geometry to enable alignment across spatial epigenomics, transcriptomics, proteomics, metabolomics, and histological data. Across 34 benchmark datasets covering 16 platforms and 6 modalities, MAPS-alignment consistently outperformed seven state-of-the-art methods, achieving average speedups exceeding 100-fold and substantially reducing GPU memory consumption. For example, it aligned 2 breast-cancer slices containing 900,000 cells in just 41 s with a peak memory of 4.8 GB, and matched manual proofreading accuracy when co-registering a mouse embryo H&E image (>1.35 million cells) with Xenium data within 76 s using less than 5 GB. Notably, it is the only method capable of aligning orthogonal modalities that lack any shared features. In such unanchored scenarios, MAPS integration accurately resolves the triple-modal anatomical architecture of the mouse cerebellum and reveals functional regionalization in the mouse tongue epithelium. Furthermore, when using paired H&E histology to predict cellular-resolution spatial multi-omics, MAPS-integration outperformed 7 competing prediction methods, improving the Pearson correlation coefficient by 10–30%.

The integrative capability of MAPS was further examined at two distinct scales. At the consortium scale, MAPS integrated 434 slices from 18 atlases, 14 platforms, and 5 molecular modalities, comprising more than 21 million cells, into a unified 3D mouse brain coordinate system. This reconstruction enabled automated cross-modal anatomical annotation via spatial-nearest-neighbors label propagation, allowing modalities devoid of transcriptomic markers, such as MALDI-MSI metabolomics and low-plex CODEX proteomics, to be mapped directly onto the Allen CCF. At the individual-laboratory scale, MAPS demonstrated a cost-effective strategy for 3D reconstruction: using routine H&E staining as a structural scaffold, it propagated dense multi-omics measurements from only 2 anchor slices through an entire tissue block of serial HCC sections (∼26 million cells).

The value of 3D reconstruction extends beyond visualization, as it also exposes systematic biases inherent in 2D analysis. In the HCC dataset, 3D Euclidean distances from different cell populations to their nearest blood vessels were consistently shorter than corresponding 2D in-plane distances. This difference reflects the fact that the nearest vascular structure may lie outside the slice in which a cell is observed. Similarly, neighborhood analysis showed that 2D measurements tended to produce stronger estimates of homotypic cell clustering, whereas volumetric neighborhoods contained a greater diversity of heterotypic cellular contacts. Together, these results demonstrate a measurable spatial truncation effect associated with conventional 2D analysis. In particular, restricting neighborhood definitions to a single slice can omit biologically relevant out-of-plane neighbors and consequently alter estimates of local cellular composition and proximity. 3D analysis therefore provides a more complete representation of tissue neighborhoods, although the biological significance of individual spatial contacts still requires interpretation in the context of cell type, interaction range, and tissue structure. Leveraging the reconstructed 3D volume, we uncovered spatially organized tumor–immune interfaces that remain entirely obscured in conventional sections. Continuous z-axial profiling revealed a depth-dependent immune state transition, characterized by a significant decline in monocyte–malignant colocalization with increasing depth, accompanied by downregulation of metabolic-stress markers (*PLIN2, NDRG1*) and upregulation of immune-regulatory genes (*LAG3, CD44*).

Several limitations also chart a course for future development. First, our reconstructions remain based on serial 2D sections with inherent registration artefacts, tissue deformations and variable sampling gaps; as true volumetric spatial profiling technologies mature, future frameworks transition from 2D slices to directly processing continuous 3D density fields and non-rigid deformations. Second, the current atlases are static snapshots, whereas biological systems evolve over time; extending toward 4D spatiotemporal representations will require models that jointly capture physical displacement, cellular state transitions and lineage trajectories. Third, integrating modalities with minimal molecular overlap remains a fundamental challenge. Beyond geometric cues, prior biological knowledge, encoded in curated databases, literature or large language models, could serve as a semantic bridge to align otherwise disconnected modalities. However, because external knowledge is inherently incomplete and biased, it should complement rather than replace directly measured spatial data.

Collectively, MAPS transforms fragmented spatial data into queryable, explorable 3D multi-modal atlases, supporting both consortium-scale reference mapping and translational discovery at the individual-laboratory level. More fundamentally, it moves the field beyond the flatland paradigm, revealing not only the limitations of 2D projections but also the deep organizational logic of tissues in 3D and uncovers biological principles, such as the spatial segregation of signaling components, that are inaccessible through 2D spatial omics alone.

## Methods

### 4.1 Datasets

#### The in-house spatial multi-omics hepatocellular carcinoma dataset

An intact hepatocellular carcinoma (HCC) tissue block was obtained from the First Affiliated Hospital of Wenzhou Medical University (Ethics approval number: 2024-R318) (see Supplementary Note 3). To facilitate downstream 3D multimodal integration, the tissue block was serially sectioned into 100 consecutive slices (N = 100). Of these, three slices (ID1, ID10, and ID100) were selected for single-cell-resolution spatial multi-omics profiling (equipped with H&E staining and imaging), while the remaining slices were subjected only to H&E staining and imaging. Regarding omics profiling, each selected slice was analyzed using Xenium Prime 5K and PhenoCycler-Fusion (PCF) for spatial transcriptomics (5,000 genes) and spatial proteomics (21 proteins), respectively (see Supplementary Note 3 for details). We then introduced a panel of interest by pooling the top 500 highly variable genes (HVGs) identified from each slice and supplementing them with canonical markers associated with tumor proliferation and immune processes, resulting in a final panel of 658 genes. Based on H&E images and histopathological characteristics, two expert pathologists manually annotated the tissue into four major regions: tumor, boundary, normal, and vascular regions.

#### Multi-modal 3D mouse brain atlas

We collected coronal sections across 18 brain atlases, covering 5 modalities and 434 slices, with a total of 21,335,013 cells/spots (Fig. 6a), including Chen et al.^94^ (Atlas 1, BARseq), Shi et al.^96^ (Atlas 2, STARmap PLUS), Zhang et al.^98^ (Atlas 3, MERFISH), Yao et al.^97^ (Atlas 4, MERSCOPE), Vizgen. (Atlas 5, MERSCOPE), 10xGenomics. (Atlas 6, Xenium), 10xGenomics. (Atlas 7, Xenium5k), 10xGenomics. (Atlas 8, Xenium), STOmics. (Atlas 9, Stereo-seq), NanoString. (Atlas 10, CosMx), Vizgen. (Atlas 11, MERFISH V2), Zeng et al.^19^ (Atlas 12, STARmap), Zeng et al.^19^ (Atlas 13, RIBOmap), Han et al.^95^ (Atlas 14, Stereo-seq), Han et al.^95^ (Atlas 15, Stereo-seq), Zhang et al.^30^ (Atlas16, CODEX), METASPACE. (Atlas17, MALDI-MSI), 10xGenomics. (Atlas18, H&E).

#### More data description

Please refer to Supplementary Note 1.

### 4.2 MAPS-alignment module

MAPS-alignment is a modality-agnostic alignment module that formulates spatial registration as a continuous optimization problem. It estimates the optimal similarity transformation (e.g., isotropic scaling, rotation, and 2D translation) solely from spatial coordinates. This geometric-only design enables seamless application across diverse modalities (see Supplementary Fig. 1). To further accommodate heterogeneous experimental scenarios, MAPS-alignment incorporates a complete pipeline that includes spatial transformation, optimization with acceleration, a coarse-to-fine initialization strategy, ROI-guided partial alignment, and multi-slice 3D reconstruction.

#### Spatial transformation

Let the spatial coordinates of the source and target slices be denoted as 2D point clouds S ∈ ℝ^*N*×2^ and T ∈ ℝ^*M*×2^, respectively, where *N* and *M* are the numbers of cells or spots. Let C_*s*_ denotes the geometric centroid of the source point cloud. MAPS applies a similarity transformation to the source coordinates, yielding the transformed point cloud S′ as:

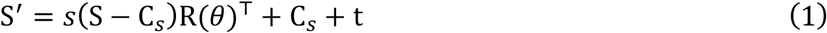

where *s* > 0 is an isotropic scaling factor, 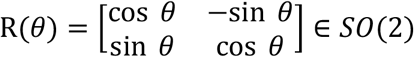 is the rotation matrix parameterized by angle *θ*, and t ∈ ℝ^2^ is the translation vector. The transformation is applied sequentially by centering the source coordinates, performing isotropic scaling and rotation, and subsequently restoring the original centroid and applying translation. Given the scaling is isotropic, the scaling and rotation operations are commutative.

#### Optimization objectives and alignment acceleration

We adopt the bidirectional Chamfer Distance^126^ as the objective function to quantify the spatial discrepancy between the transformed source S′ and the target T:

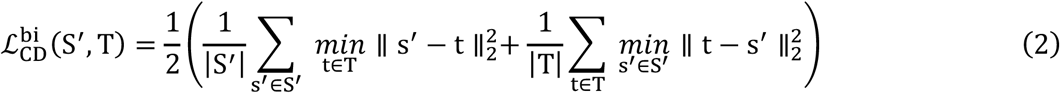

For large-scale spatial omics datasets, computing the pairwise distance matrix across the whole slices can be computationally prohibitive and easily exceed available GPU memory^38,43,84,127^.

To circumvent this bottleneck, we introduce a stochastic subsampling strategy. Specifically, at each optimization epoch, *K* points are randomly sampled from both the transformed source and target point clouds, and the alignment loss is computed over sampled subsets. This approach provides two key advantages. It enormously reduces peak GPU memory, enabling alignment for extremely large-scale datasets. Moreover, stochastic subsampling introduces variability into the optimization process, analogous to mini-batch stochastic gradient descent, reducing sensitivity to unfavorable local minima.

#### Alignment initialization strategy

To mitigate the sensitivity towards non-convex geometric alignment initialization^128,129^, we implement a coarse-to-fine optimization strategy. The isotropic scaling factor is first initialized heuristically based on the ratio of the mean distances of points to their respective geometric centroids, while the translation vector is initialized according to the displacement between the source and target centroids. A coarse rotation search is then performed over 180 uniformly spaced angles within [0,2*π*), with the Chamfer loss evaluated on a random subset of points to identify the optimal initial rotation. Finally, all transformation parameters are jointly refined via gradient-based optimization using Adam optimizer (betas = (0.9, 0.999)).

#### ROI-guided partial alignment

To address scenarios in which the source slice overlaps only partially with the target one, rendering global centroid-based initialization unreliable, we design an ROI-guided partial alignment module. Specifically, a sliding square window, with its size determined by the bounding-box dimensions of the source point cloud, is scanned over the target slice with a stride equal to 3% of the window size to generate candidate ROIs. Windows containing fewer than 50 points are excluded as background. For each valid ROI, its local centroid is computed, and 500 points are randomly subsampled for efficient initialization.

For every valid ROI, a parallel coarse rotation search is then performed using the centered and subsampled source point cloud. At each candidate angle, the unidirectional Chamfer distance from the rotated source to the ROI is computed, while the corresponding translation is initialized as the displacement between the ROI and source centroids. The ROI-angle pair that minimizes this loss is selected, providing an initial rotation and translation anchor, which are subsequently used to initialize Adam-based fine-tuning on the full point clouds, ensuring both robust partial alignment and global consistency.

#### Multi-slice alignment and 3D reconstruction

MAPS provides two alignment modes for multi-slices datasets: reference mode and sequential mode. Given a collection of AnnData objects representing multiple tissue slices, reference mode aligns all slices directly to a user-specified reference slice. In sequential mode, the first slice is kept fixed, and each subsequent slice is aligned to its immediately preceding slice (i.e., *i*-th slice aligned to (*i*−1)-th). The resulting transformations are propagated across consecutive slices, enabling reconstruction of a spatially coherent 3D tissue volume.

### 4.3 MAPS-integration-U module

MAPS-integration-U is a geometry-aware diagonal integration module that integrates unpaired spatial data with no shared features into a unified latent space. Specifically, following MAPS alignment, the integration pipeline constructs both intra- and inter-slice spatial graphs and applies a local pooling network^84^ for diagonal feature mapping, thereby establishing a unified latent representation. By exploiting spatial correspondences across slices, this strategy encourages feature consistency across spatially matched regions while preserving modality-specific information and promoting cross-modality alignment in the shared latent space (see Supplementary Fig. 1).

#### Spatial graph construction

For each slice, we follow SpaLP^84^ to use a KD-Tree algorithm to retrieve the *K*_intra_ spatial nearest neighbors for each cell, thereby constructing a spatial graph. The neighbor indices are stored as an index matrix 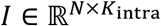. For inter-slice graph construction, we establish a cross-slice neighbor index that maps points from one modality to their nearest neighbors in the coordinate space of another modality. We construct a KD-Tree over the target spatial coordinates 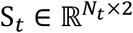 and query each source coordinates 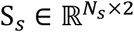 for its *K* nearest neighbors in the target slice, yielding a cross-slice index matrix 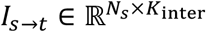. In three-modal setting, we construct two index matrixes, I_*modal*1→*modal*3_ and *I*_*modal*2 →*modal*3_ to bridge modality 1 and modality 2 with modality 3, respectively.

#### Cross-modal diagonal integration

We employ two local pooling encoders together with modality-specific linear decoders to learn intra- and inter-modality features. For intra-modality features 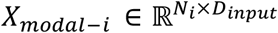, we employ Multi-Layer Perceptron (MLP) to extract the initial representation and further construct a neighborhood feature matrix *L*_*i*_:

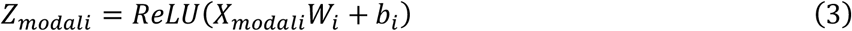

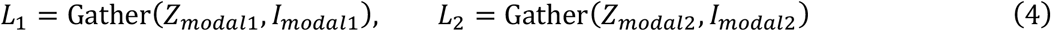

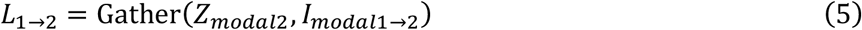

where 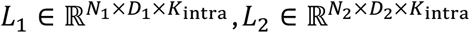 and 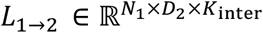. Gather represents extracting neighbor features from *Z* by index *I*. We then perform local pooling operation to aggregate the *K* neighboring features into a compact latent representation for each cell. Specifically, for a given neighborhood feature matrix *L*, we first compute channel attention scores across the neighbor dimension:

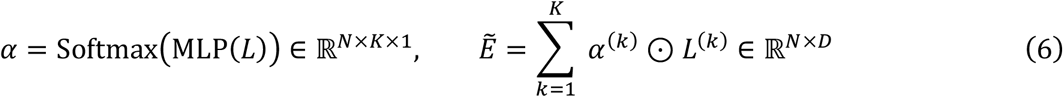

where ⊙ denotes element-wise multiplication. The aggregated features are then passed through a linear projection layer to obtain the final modality-specific embeddings. Consequently, we derive the intra-modality embeddings 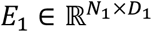 and 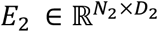, along with the cross-modality translated embedding 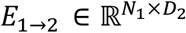. To preserve modality-specific information, we employ two independent linear decoders *g*_1_ and *g*_2_ to reconstruct the original feature from the embeddings:

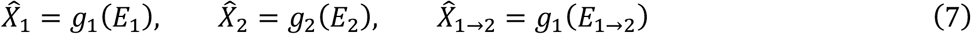

where 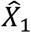 is reconstructed from the intra-modal embedding of modality 1, 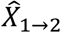 is derived from the cross-modal embedding translated from modality 2. Notably, with 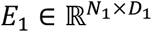 and 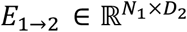 sharing the same latent dimension *D*.

#### Model training

To facilitate biologically meaningful cross-modal integration, we impose two complementary constraints during model training: (i) a reconstruction loss, which encourages the learned embeddings to preserve modality-specific information by minimizing the Mean Squared Error (MSE) between the input and reconstructed features; (ii) a cross-modal alignment loss, which minimizes the representation discrepancy between the native modality 1 embedding *E*_1_ and translated embedding *E*_1→2_ derived from modality 2:

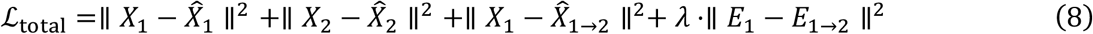

where *λ* is a hyperparameter controlling the alignment strength. By jointly optimizing the two terms, the model learns a shared latent space in which representations from different modalities are explicitly aligned while retaining modality-specific information. For scenarios involving three modalities, we designate Modality 3 as the anchor and construct two cross-modal mappings from Modality 1 and Modality 2 to Modality 3:

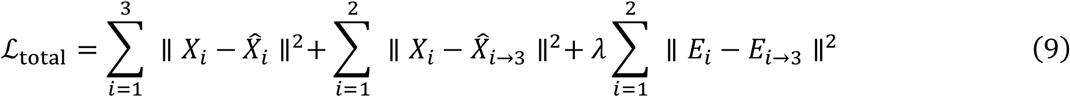

### 4.4 MAPS-integration-P module

MAPS-integration-P aims to propagate multi-omics molecular information across the entire tissue block using morphology features from H&E images. Specifically, the neural network is trained using both omics-profiled slices and their adjacent, unprofiled slices. The profiled slices are used to learn the correspondence between tissue morphology and molecular features, whereas the adjacent slices are incorporated to mitigate inter-slice domain shifts and improve the robustness of molecular inference across the tissue volume. The detailed training procedure is described in the following sections (see Supplementary Fig. 1).

#### Preliminary feature extraction

After having the sequenced multi-omics slices and their adjacent imaging-only slices, we first extract the cellular-level morphology features using pre-trained histology foundation models (Supplementary Note 7). Any given cell in sequenced and imaging-only slices can be represented as *x*^∗^ = (*f,y*_1_ … *y*_*m*_) and *x*^ = (*f*), respectively, where *f* denotes the morphology features comprising both global (*f*_global_) and local (*f*_local_) information, and *y* indicates the omics features across *m* categories.

We first employ two Non-Linear Blocks (NLB) to independently recalibrate the global (*f*_global_) and local (*f*_local_) components. These refined features are then concatenated into a unified multi-scale representation:

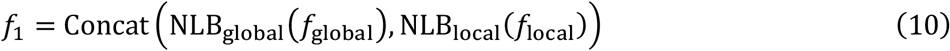

Each NLB comprises two linear layers, each followed by a LeakyReLU activation function. To align these morphological features with the underlying biological data, *m* joint non-linear blocks (corresponding to the *m* omics types), process *f*_1_ to generate omics-specific morphology features. Each NLB_joint_ comprises a single linear layer followed by a LeakyReLU activation. The multi-omics representation is defined as the stack of these outputs:

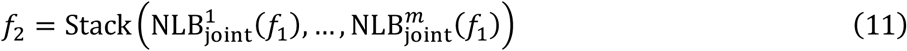

where *f*_2_ ∈ ℝ^*m*×*d*^.

#### ProtoEncoder

To facilitate high-fidelity feature extraction across heterogeneous biological modalities, the integrated multi-omics features are processed through a novel ProtoEncoder. Architecturally inspired by the BEiT (Bidirectional Encoder representation from Image Transformers)^130^, this module leverages learnable prototypes as latent anchors to capture complex semantic abstractions. We denote the set of learnable prototypes as *P* = [*p*_1_, …, *p*_*m*_], i.e., one prototype for one omics layer. The *i*_th prototype is denoted as 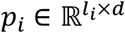, with *l*_*i*_ and *d* indicating the omics and feature dimensions, respectively. These prototypes are initialized as trainable parameters, allowing the model to learn a discrete latent codebook automatically. After this, *f*_2_ and *P* were passed through the cross-attention computation.

#### Cross-Attention Computation

To integrate the image features (*f*_2_) and the prototype set *P*, we employ a Multi-Head Cross-Attention (MHCA) mechanism followed by a Feed-Forward Network (FFN)^131^. The interaction is modeled as a query-key-value operation, i.e., query is from *f*_2_, and key and value are from *P*. The single-head scaled dot-product attention can be formulated as follows:

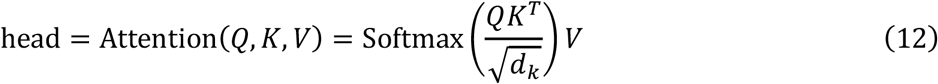

where 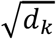 is the scaling factor. The implemented Multi-head Cross-attention allows the model to jointly attend to information from disparate feature subspaces:

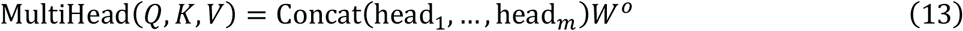

To ensure stability and capture non-linearities, the output of the MHCA is implemented under the pre-layer normalization (Pre-LN)^132^ strategy:

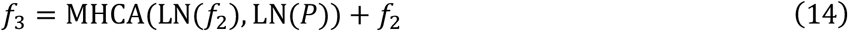

where LN represents the layer normalization operation. In this context, the MHCA allows the query (*f*_2_) to “attend” to the most informative components of the prototypes, effectively mapping the input data into a refined omics-informed representation. Following MHCA, features are further processed by a Pre-LN FFN^132^ to capture complex non-linearities, as follows:

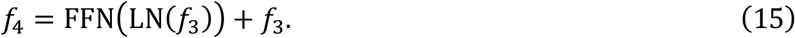

#### Self-attention Computation

Immediately after the cross-attention stage, the refined features undergo a self-attention computation to establish global feature dependencies. While the underlying architecture of the MHSA and its associated FFN remains consistent with the cross-attention block, the query, key, and value (*Q, K, V*) are now all derived from the feature set itself, i.e., *f*_4_ . We repeat the above computation process (Cross-attention & Self-attention) for two times and denoted the final generated features as *f*_latent_ ∈ ℝ^*m*×*d*^ (latent embedding).

#### Omics decoder

Distinct from previous methods that utilize a linear regression layer to predict the omics expression values, our framework utilizes the learned prototypes as a more robust alternative. Initially, we employed a non-linear block (identical structure as previous ones) for feature recalibration as omics embeddings *e* ∈ ℝ^*m*×*d*^. Afterwards, we chunked the learned features along the omics layer dimension *e* = [*e*_1_, …, *e*_*m*_]. Then, the learned features were multiplied by their corresponding prototypes to regress the omics expression values:

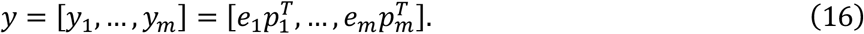

#### Adaptive branch to remove stain variations

As illustrated in Supplementary Fig. 20, the staining process brings a significant domain shift in the feature space. Directly deploying a trained model on the sequenced slices to the imaging-only slices could result in unstable and inaccurate prediction due to the domain gap. To enhance generalization, we incorporate a Gradient reversal layer (GRL)^133^, a widely adopted domain adaptation technique. Mathematically, the GRL serves as a critical bridge between the extracted features and a domain discriminator. The GRL is a specialized operator that leaves the input unchanged during the forward pass, ensuring that the feature representations are propagated to the discriminator for classification. However, during the backward pass (backpropagation), the GRL scales the gradients by a negative hyperparameter −*λ*:

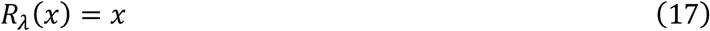

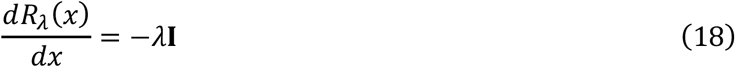

where **I** is the identity matrix and *x* simply represents any input features. This inversion compels the network to update its weights in a direction that maximizes the discriminator’s loss, thereby pushing the encoder to learn features invariant to domain-specific noise or batch effects.

In this study, the adaptive branch comprises a gradient reversal layer and a discriminator. Here, the discriminator comprises two linear layers with the first one followed by a LeakyReLU activation function. We calculated the cross-entropy loss for the discriminator between slide ID and the prediction. In this study, there are in total two adaptive branches accommodating *f*_2_ and *f*_latent_.

#### Loss Function Formulation

Apart from the loss functions utilized in the gradient reversal layer for domain adaptation, we also have two varieties of loss functions: weighted mean squared error loss and mean square error loss. The mean square error loss can be formulated as follows:

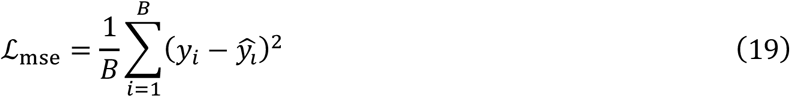

where *B* is the total number of cells in each mini-batch and *y*_*i*_ and 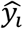 indicate the measured and predicted expressions, respectively. In terms of the weighted one, we simply utilized the exponential value of the omics features as weight:

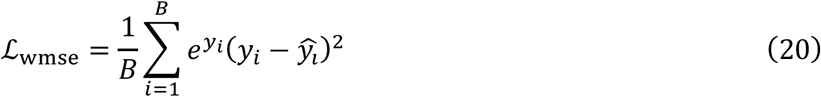

In this study, we utilize ℒ_mse_ for the spatial proteomic prediction. Given the inherent sparsity of spatial transcriptomics data, we introduced the weighted one ℒ_wmse_ for computation.

#### Cell type inference

Characterizing cellular identities is fundamental to deciphering tissue architecture; however, transferring established cell-type labels from sequenced reference data to imaging-only slices presents a substantial computational challenge. To address this, we developed a robust retrieval-based strategy that leverages the high-fidelity latent embeddings learned by MAPS-integration-P. Specifically, we constructed a reference library using the latent embeddings 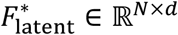 and corresponding cell type labels Ct^∗^ of cells from the sequenced slices. For each cell in an imaging-only slice, we computed the embedding-space distances between its latent representation and all cells in the reference library. The top- *k* nearest reference cells were then identified, and cell type annotations were transferred through majority voting. A cell-type label is assigned only if a single type constitutes a strict consensus (exceeding *k*/2) among the top-*k* neighbors; otherwise, the cell was considered insufficiently confident and excluded from downstream analysis.

### 4.5 Data preprocessing

#### MAPS-alignment

For omics-to-omics alignment, spatial coordinates were extracted from obsm[‘spatial’] of the source and target AnnData^134^ objects. For alignment involving imaging data (omics-to-image or image-to-image), Cellpose^135^ was employed to automatically segment H&E or immunofluorescence (IF) images (Supplementary Note 6), and the central coordinates of the segmented cells were utilized for subsequent computing.

#### MAPS-integration-U

For transcriptomics and metabolomic data, expression counts were library-size normalized and log-transformed via SCANPY^136^. Afterwards, the top 2,000 highly variable features were selected as input to the encoder. For proteomics data, we applied centered log ratio normalization to the raw protein expression counts. For image data, the pre-trained histopathology foundation model UNI^137^ was used to extract the image representations, followed by L2-normalization (Supplementary Note 7).

#### MAPS-integration-P

For both transcriptomic and proteomic data, expression counts were log-transformed using SCANPY, followed by max-min normalization. For image data, we applied the pre-trained histopathology foundation model UNI^137^ to extract the image representation (Supplementary Note 7).

#### More benchmark data preprocessing

Please refer to Supplementary Note 5.

### 4.6 MAPS-Explorer module

MAPS-Explorer is a self-hosted platform for interactive 2D and 3D visualization and analysis of spatial omics data. It supports deployment across major operating systems and device types without requiring external database or object-storage services.

#### System architecture and data storage

MAPS-Explorer consists of a Python backend implemented in Django 4.2, a browser-based frontend implemented in JavaScript and WebGL, and a custom binary container termed MBIN. The backend provides RESTful APIs, background task execution, and project-level metadata management, whereas the frontend performs interactive rendering and analysis. An optional read-only mode restricts write operations for publicly hosted projects. MBIN stores sparse expression matrices and associated metadata in independently indexed, z lib-compressed blocks. Each block records its data type, location, and byte length, allowing individual genes or metadata columns to be retrieved without loading the complete dataset. Frequently accessed project metadata, including file descriptors, observation columns, and gene lists, are cached locally in SQLite to avoid repeated deserialization of source H5AD files.

#### GPU-based 3D rendering

The MAPS-Explorer 3D Viewer renders large point clouds using WebGL. Spatial coordinates and annotations are transferred from the backend as byte buffers, and coordinate data are uploaded to the client-side GPU for subsequent rendering and transformation. Previously accessed metadata are retained in an LRU cache to reduce repeated data transfer. Rendering is event driven and is triggered only when the scene changes. Camera transformations are applied to GPU-resident coordinates rather than recomputing point positions on the CPU. Spatial clipping along the x-, y-, and z-axes is also performed on the GPU. Metadata filtering retains the original point geometry and updates display attributes, avoiding repeated reconstruction and upload of the point cloud.

#### Voxelized surface reconstruction

MAPS-Explorer provides voxelized surface reconstruction as a complementary representation to discrete point clouds. Mesh generation is performed in an independent Web Worker using a workflow adapted from the Marching Cubes algorithm^138^, with additional processing to accommodate cavities, inter-section discontinuities, and incomplete boundaries in sparse spatial point clouds. The reconstructed mesh is returned to the main thread for GPU rendering and can be displayed together with the underlying point cloud. 3D meshes can also be exported in GLB format.

#### MAPS-Explorer surface dynamics

Surface dynamics analyzes 3D Label Shell meshes generated from spatial point clouds. It provides geometric statistics, molecular signal mapping, surface-normal gradients, and region-restricted surface analysis.

#### Surface geometry

Connected mesh components are identified by shared vertices after coordinate quantization to 10^-4^. Components containing fewer than two triangles are discarded. For triangle t with vertices a, b, and c, its area is

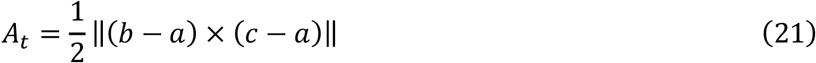

total surface area is *A* = ∑_*t*_ *A*_*t*_, and enclosed volume is estimated as 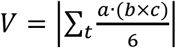. Where, A and V are expressed in the normalized coordinate system. Volume estimates assume a closed mesh with consistent triangle orientation. Cell coverage is approximated as the number of points within the axis-aligned bounding box of each mesh component and should therefore not be interpreted as an exact point-in-polyhedron count.

#### Surface molecular signal

Surface points are sampled approximately uniformly by weighting triangles according to area. The number of sampling points *N*_*s*_ ranges from 1,000 to 5,000. The local neighborhood radius is estimated from both surface area and point-cloud volume:

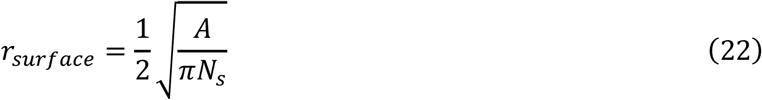

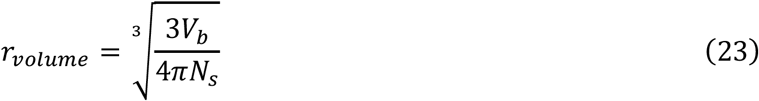

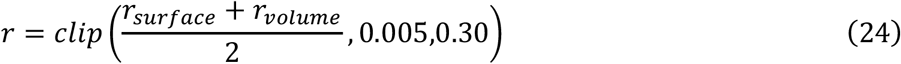

where A is total surface area, *N*_*s*_ is the number of surface samples, and *V*_*b*_ is the point-cloud bounding-box volume.

For surface sample s, nearby cells are selected within a cylindrical neighborhood oriented along the local surface normal. The mapped molecular signal is

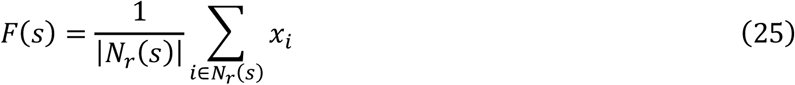

where *N*_*r*_(*s*) denotes cells within radius r around the sampled surface position and *x*_*i*_ is the corresponding gene-expression or annotation-derived signal. 3D spatial hashing is used to accelerate neighborhood queries.

#### MAPS-Explorer visualization

MAPS-Explorer provides three modes for visualizing aligned spatial data: Global Align, Ref Align, and Impute Align. Spatial registration itself is performed by MAPS or another compatible procedure, and input H5AD files therefore contain precomputed aligned x- and y-coordinates. **(1) Global Align**. For N aligned sections, the third coordinate z is assigned from a user-selected numerical section variable. Categorical annotations are encoded using a shared label vocabulary and stored independently by metadata column. Gene-expression vectors are similarly indexed by gene, enabling selective retrieval without reparsing the original H5AD files. **(2) Ref Align**. Ref Align uses the same coordinate and metadata representation but restricts visualization to a target section and its designated reference section. **(3) Impute Align**. Impute Align inserts a newly aligned section into an existing three-dimensional reference atlas. The insertion position can be specified manually or obtained from the MAPS atlas-matching result stored in uns.atlas_match.best_atlas_slice_idx. Inserted data are maintained separately from the reference atlas and concatenated at visualization time, leaving the original atlas unchanged.

#### MAPS-Explorer cell density viewer

The Cell Density Viewer compares the projected spatial enrichment of categorical cell populations. Visible cells are projected onto a selected two-dimensional plane, binned independently by label, Gaussian smoothed, and rendered as three-dimensional density surfaces. Only cells passing the current label visibility, file visibility, metadata filters, and three-dimensional slab filters are included. In 3D mode, the x-, y-, or z-axis can be selected as the collapse dimension before density estimation. Projected cells are assigned to an *n* × *n* grid, where 20 ≤ *n* ≤ 100 and the default is *n* = 80. For cell label c,

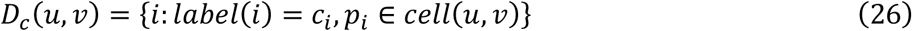

where *D*_*c*_(*u, v*) is the cell count in grid bin (*u, v*). Each grid is smoothed using a separable Gaussian kernel,

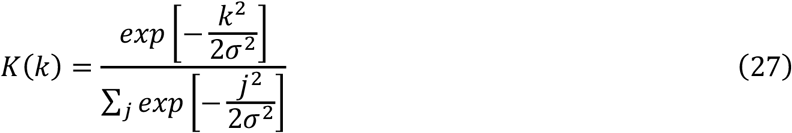

with kernel radius ⌈3σ⌉. Smoothed densities are 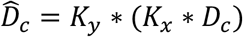, where σ controls the smoothing strength. To retain comparable height scales across cell populations, all labels share a global maximum 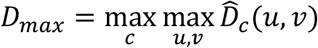 and the displayed surface height is

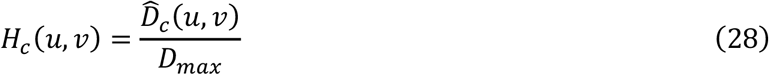

Thus, differences in surface height reflect differences in absolute smoothed cell counts within the same view rather than independent normalization of each population. The resulting grids are triangulated and rendered by WebGL. Density calculation, smoothing, and mesh construction are executed in a Web Worker. Because density is calculated after projection, the output represents smoothed projected cell counts rather than a physical density in cells per unit area; active three-dimensional slab filters can be used to restrict the depth range before projection.

#### Gene co-expression visualization module

Please reference supplementary note 8.

### 4.7 Data perturbation for alignment benchmark

For each dataset, we designated one slice as the fixed reference, while randomly perturbing the remaining slices by sampling a rotation angle from a uniform distribution over [0, 2π) and applying translations along the x- and y-axes based on each slice’s coordinate range. This procedure was repeated independently five times per dataset.

### 4.8 Spatial domain identification

Since MAPS-integration-U produces spatially resolved embeddings, tissue spatial domains can be directly identified using KMeans clustering implemented in the scikit-learn package^139^. In contrast, MAPS-integration-P produces the cellular representation without explicitly incorporating spatial context. Therefore, CellCharter^140^ was applied to the learned cellular representations *f*_latent_ to incorporate spatial neighborhood information for subsequent domain identification.

### 4.9 PTR analysis

To characterize the cellular-level relative abundance between translated proteins and their coding genes, we referred to CAST^40^ and defined a Protein-to-Transcript Ratio (PTR) score. For each cell *j* and each matched omics pair *i*, protein and RNA expression matrices were independently normalized by library size scaling to a fixed total of 10,000 counts per cell:

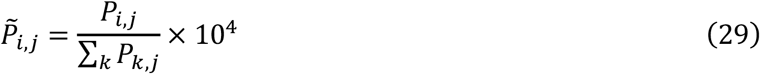

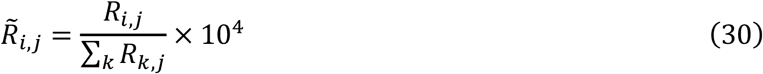

where 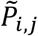 and 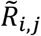 denote the protein and RNA counts, respectively, *k* is the number of paired genes and proteins. We then defined the raw protein-to-transcript ratio (rPTR) as the log2-transformed ratio:

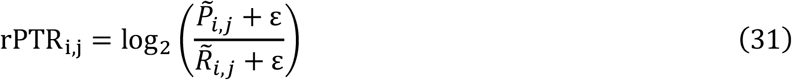

where ε = 10^−10^ was introduced to avoid numerical instability due to zero values. Because different genes exhibit substantial variation in overall abundance and dynamic range, direct comparison of raw PTR values may be confounded by gene-specific distributions. Therefore, for each matched omics pair, raw PTR values were standardized across all cells:

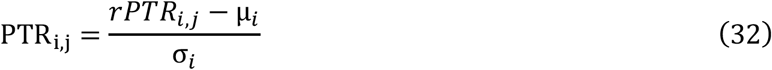

where *μ*_*i*_ and *σ*_*i*_ represent the mean and standard deviation of rPTR across all cells for matched pair *i*.

The standardized PTR reflects the relative protein-to-transcript ratio of each cell, with positive and negative values indicating above- and below-average ratios, respectively. As the true translation rate cannot be directly measured, PTR should be interpreted as a relative indicator of potential post-transcriptional variation rather than an absolute biological signal.

### Benchmarking methods and evaluation metrics

We compared MAPS-alignment with 7 state-of-the-art alignment methods: PASTE^38^, STalign^51^, SLAT^52^, SANTO^53^, CAST^54^, GPSA^42^ and STAIR^55^. We compared MAPS-integration-U with SpatialMETA^59^ on mouse cerebellum. We compared MAPS-integration-P with 7 state-of-the-art H&E -based predictive methods: SpatialEx^49^, GHIST^77^, OmiCLIP^78^, DeepPT^79^, iStar^80^, mclSTExp^81^, ST-Net^82^. All of benchmarking methods were executed based on their provided vignettes. Detailed evaluation metrics are available at Supplementary Note 9.

### Software and package versions

All analyses were performed in python (3.10.13) using the following package versions: scanpy (1.9.8), torch (2.2.0), scipy (1.12.0), pandas (2.2.3), numpy (1.26.3), scikit-learn (1.4.0), numba (0.59.0), llvmlite (0.42.0), anndata (0.10.5.post1), leidenalg (0.10.2), igraph (0.11.8), scikit-misc (0.5.1), pynndescent (0.5.13), einops (0.8.2) and tifffile (2025.5.10).

## Code availability

The python package source code of MAPS is available at https://github.com/dbjzs/MAPS. The jupyter notebook tutorials for reproducing main results in this paper and MAPS-Explorer tutorials are available at https://mapspatial.readthedocs.io/en/latest/. The MAPS-Explorer interactive website and demo video are available at https://bioinfor.imu.edu.cn/maps-explorer/.

## Data availability

All benchmarking data and results are provided in this paper. All h5ad files have been uploaded and are freely available at https://zenodo.org/records/22100091. The in-house multi-modal HCC dataset will be publicly available after publication.

## Acknowledgements

This research was funded by the National Natural Scientific Foundation of China (No. 62571279 (Y.Z.), No. 62303119 (Z.Y.), No. 32470706 (Z.Y.)), the Group Project of Developing Inner Mongolia through Talents (No. 2025TEL25 (Y.Z.)), the Computational Biology Program (No. 25JS2850200 (Z.Y.)) of Science and Technology Commission of Shanghai Municipality (STCSM), the Central Guidance Fund for Local Science and Technology Development (No. 2024ZY0168 (Y.Z.)), Shanghai Municipal Science and Technology Major Project (2023SHZDZX02 (B.Q.)), the National Key R&D Project of China (2023YFC3402501 and 2023YFC3402500 (B.Q.)) and the AI for Science Foundation of Fudan University (FudanX24A1031 (B.Q.)). We thank Jianmin Li (Department of Pathology, The First Affiliated Hospital of Wenzhou Medical University, Wenzhou, China) and Xiaoping Yi (Chongqing University Three Gorges Hospital) for their pathological annotations.

## Author Contributions Statement

Z.Y., B.Q., Y.Z. and Z.W. conceptualized and supervised the project. B.D., Y.Y., Z.W. and Y. L. designed the computational method, contributed to the analysis of data. B.D., Y.Y., Z.W., Y. L., X.Y., C.W. and Z.Y. prepared the figures and tables. B.D., Y.Y., Z.W., Y. L., and P.H. prepared the tutorials. B.D., Y.Y., Z.W., Y. L., C.S., J.H., X.Z., H.C., D.Z., Q.Z., Y.D., Z.H., Y.X., G.C., Z.F., and B.Q. collected and processed the datasets. B.D., Y.Y., Z.W., Y. L., S.L., J.F., S.X., W.H., Y.Z. and Z.Y. wrote the manuscript. All authors have read, revised, and approved the final manuscript.

## References

1 Gulati, G. S., D’Silva, J. P., Liu, Y., Wang, L. & Newman, A. M. Profiling cell identity and tissue architecture with single-cell and spatial transcriptomics. Nature Reviews Molecular Cell Biology 26, 11–31 (2025). 10.1038/s41580-024-00768-2

2 Rao, A., Barkley, D., França, G. S. & Yanai, I. Exploring tissue architecture using spatial transcriptomics. Nature596, 211–220 (2021). 10.1038/s41586-021-03634-9

3 Longo, S. K., Guo, M. G., Ji, A. L. & Khavari, P. A. Integrating single-cell and spatial transcriptomics to elucidate intercellular tissue dynamics. Nature Reviews Genetics 22, 627–644 (2021). 10.1038/s41576-021-00370-8

4 Moffitt, J. R., Lundberg, E. & Heyn, H. The emerging landscape of spatial profiling technologies. Nature Reviews Genetics 23, 741–759 (2022). 10.1038/s41576-022-00515-3

5 Jain, S. & Eadon, M. T. Spatial transcriptomics in health and disease. Nature Reviews Nephrology 20, 659–671 (2024). 10.1038/s41581-024-00841-1

6 Chen, A. et al. Spatiotemporal transcriptomic atlas of mouse organogenesis using DNA nanoball-patterned arrays.Cell 185, 1777–1792 e1721 (2022). 10.1016/j.cell.2022.04.003

7 Chen, K. H., Boettiger, A. N., Moffitt, J. R., Wang, S. & Zhuang, X. RNA imaging. Spatially resolved, highly multiplexed RNA profiling in single cells. Science 348, aaa6090 (2015). 10.1126/science.aaa6090

8 He, S. et al. High-plex imaging of RNA and proteins at subcellular resolution in fixed tissue by spatial molecular imaging. Nat Biotechnol 40, 1794–1806 (2022). 10.1038/s41587-022-01483-z

9 Janesick, A. et al. High resolution mapping of the tumor microenvironment using integrated single-cell, spatial and in situ analysis. Nat Commun 14, 8353 (2023). 10.1038/s41467-023-43458-x

10 Lee, Y. et al. XYZeq: Spatially resolved single-cell RNA sequencing reveals expression heterogeneity in the tumor microenvironment. Sci Adv 7 (2021). 10.1126/sciadv.abg4755

11 Lubeck, E., Coskun, A. F., Zhiyentayev, T., Ahmad, M. & Cai, L. Single-cell in situ RNA profiling by sequential hybridization. Nat Methods 11, 360–361 (2014). 10.1038/nmeth.2892

12 Moses, L. & Pachter, L. Museum of spatial transcriptomics. Nat Methods 19, 534–546 (2022). 10.1038/s41592-022-01409-2

13 Oliveira, M. F. et al. High-definition spatial transcriptomic profiling of immune cell populations in colorectal cancer. Nat Genet 57, 1512–1523 (2025). 10.1038/s41588-025-02193-3

14 Rao, A., Barkley, D., Franca, G. S. & Yanai, I. Exploring tissue architecture using spatial transcriptomics. Nature 596, 211–220 (2021). 10.1038/s41586-021-03634-9

15 Rodriques, S. G. et al. Slide-seq: A scalable technology for measuring genome-wide expression at high spatial resolution. Science 363, 1463–1467 (2019). 10.1126/science.aaw1219

16 Tian, L., Chen, F. & Macosko, E. Z. The expanding vistas of spatial transcriptomics. Nat Biotechnol 41, 773–782 (2023). 10.1038/s41587-022-01448-2

17 Wang, X. et al. Three-dimensional intact-tissue sequencing of single-cell transcriptional states. Science 361 (2018). 10.1126/science.aat5691

18 Sui, X. et al. Scalable spatial single-cell transcriptomics and translatomics in 3D thick tissue blocks. Nat Methods22, 2574–2584 (2025). 10.1038/s41592-025-02867-0

19 Zeng, H. et al. Spatially resolved single-cell translatomics at molecular resolution. Science 380, eadd3067 (2023). 10.1126/science.add3067

20 Buchberger, A. R., DeLaney, K., Johnson, J. & Li, L. Mass Spectrometry Imaging: A Review of Emerging Advancements and Future Insights. Anal Chem 90, 240–265 (2018). 10.1021/acs.analchem.7b04733

21 Caprioli, R. M., Farmer, T. B. & Gile, J. Molecular imaging of biological samples: localization of peptides and proteins using MALDI-TOF MS. Anal Chem 69, 4751–4760 (1997). 10.1021/ac970888i

22 Ma, X. & Fernandez, F. M. Advances in mass spectrometry imaging for spatial cancer metabolomics. Mass Spectrom Rev 43, 235–268 (2024). 10.1002/mas.21804

23 Wiseman, J. M., Ifa, D. R., Song, Q. & Cooks, R. G. Tissue imaging at atmospheric pressure using desorption electrospray ionization (DESI) mass spectrometry. Angew Chem Int Ed Engl 45, 7188–7192 (2006). 10.1002/anie.200602449

24 Goltsev, Y. et al. Deep Profiling of Mouse Splenic Architecture with CODEX Multiplexed Imaging. Cell 174, 968–981 e915 (2018). 10.1016/j.cell.2018.07.010

25 Hu, B. et al. High-resolution spatially resolved proteomics of complex tissues based on microfluidics and transfer learning. Cell 188, 734–748 e722 (2025). 10.1016/j.cell.2024.12.023

26 Liao, S. et al. Integrated Spatial Transcriptomic and Proteomic Analysis of Fresh Frozen Tissue Based on Stereo-seq. bioRxiv, 2023.2004.2028.538364 (2023). 10.1101/2023.04.28.538364

27 Deng, Y. et al. Spatial profiling of chromatin accessibility in mouse and human tissues. Nature 609, 375–383 (2022). 10.1038/s41586-022-05094-1

28 Llorens-Bobadilla, E. et al. Solid-phase capture and profiling of open chromatin by spatial ATAC. Nature Biotechnology 41, 1085–1088 (2023). 10.1038/s41587-022-01603-9

29 Zhang, D. et al. Spatial epigenome-transcriptome co-profiling of mammalian tissues. Nature 616, 113–122 (2023). 10.1038/s41586-023-05795-1

30 Zhang, D. et al. Spatial dynamics of brain development and neuroinflammation. Nature 647, 213–227 (2025). 10.1038/s41586-025-09663-y

31 Zhu, M. et al. Super-CUT&Tag: A Sensitive, Spatially Resolved Approach for Epigenomic Profiling in Tissue Sections. bioRxiv, 2025.2012.2008.692886 (2025). 10.64898/2025.12.08.692886

32 Network, B. I. C. C. A multimodal cell census and atlas of the mammalian primary motor cortex. Nature 598, 86–102 (2021). 10.1038/s41586-021-03950-0

33 Regev, A. et al. The Human Cell Atlas. eLife 6, e27041 (2017). 10.7554/eLife.27041

34 Rozenblatt-Rosen, O. et al. The Human Tumor Atlas Network: Charting Tumor Transitions across Space and Time at Single-Cell Resolution. Cell 181, 236–249 (2020). 10.1016/j.cell.2020.03.053

35 Yao, Z. et al. A high-resolution transcriptomic and spatial atlas of cell types in the whole mouse brain. Nature 624, 317–332 (2023). 10.1038/s41586-023-06812-z

36 Yayon, N. et al. A spatial human thymus cell atlas mapped to a continuous tissue axis. Nature 635, 708–718 (2024). 10.1038/s41586-024-07944-6

37 Liu, M. et al. 3D multi-omics tumour atlases: from technology to biology and clinical translation. Nature Reviews Cancer (2026). 10.1038/s41568-026-00940-0

38 Zeira, R., Land, M., Strzalkowski, A. & Raphael, B. J. Alignment and integration of spatial transcriptomics data.Nat Methods 19, 567–575 (2022). 10.1038/s41592-022-01459-6

39 Qiu, X. et al. Spatiotemporal modeling of molecular holograms. Cell 188, 1744 (2025). 10.1016/j.cell.2025.02.026

40 Tang, Z. et al. Search and match across spatial omics samples at single-cell resolution. Nat Methods 21, 1818–1829 (2024). 10.1038/s41592-024-02410-7

41 Dai, B. et al. 3d-OT: a deep geometry-aware framework for heterogeneous slices alignment of spatial multi-omics.Nat Methods (2026). 10.1038/s41592-026-03034-9

42 Jones, A., Townes, F. W., Li, D. & Engelhardt, B. E. Alignment of spatial genomics data using deep Gaussian processes. Nature Methods 20, 1379–1387 (2023). 10.1038/s41592-023-01972-2

43 Yan, Y. et al. Benchmarking alignment methods for spatial transcriptomics data. Nat Comput Sci 6, 524–541 (2026). 10.1038/s43588-026-00977-z

44 Birk, S. et al. Quantitative characterization of cell niches in spatially resolved omics data. Nat Genet 57, 897–909 (2025). 10.1038/s41588-025-02120-6

45 He, Y. et al. Towards a universal spatial molecular atlas of the mouse brain. bioRxiv, 2024.2005.2027.594872 (2024). 10.1101/2024.05.27.594872

46 Coleman, K. et al. Resolving tissue complexity by multimodal spatial omics modeling with MISO. Nat Methods22, 530–538 (2025). 10.1038/s41592-024-02574-2

47 Long, Y. et al. Deciphering spatial domains from spatial multi-omics with SpatialGlue. Nat Methods 21, 1658–1667 (2024). 10.1038/s41592-024-02316-4

48 Yan, X. et al. Mosaic integration of spatial multi-omics with SpaMosaic.Nat Genet 58, 1126–1137 (2026). 10.1038/s41588-026-02573-3

49 Liu, Y. et al. High-parameter spatial multi-omics through histology-anchored integration. Nat Methods 23, 373–386 (2026). 10.1038/s41592-025-02926-6

50 Zeira, R., Land, M., Strzalkowski, A. & Raphael, B. J. Alignment and integration of spatial transcriptomics data.Nature Methods 19, 567–575 (2022). 10.1038/s41592-022-01459-6

51 Clifton, K. et al. STalign: Alignment of spatial transcriptomics data using diffeomorphic metric mapping. Nature Communications 14, 8123 (2023). 10.1038/s41467-023-43915-7

52 Xia, C.-R., Cao, Z.-J.Tu, X.-M. & Gao, G. Spatial-linked alignment tool (SLAT) for aligning heterogenous slices.Nature Communications 14, 7236 (2023). 10.1038/s41467-023-43105-5

53 Li, H. et al. SANTO: a coarse-to-fine alignment and stitching method for spatial omics. Nature Communications15, 6048 (2024). 10.1038/s41467-024-50308-x

54 Tang, Z. et al. Search and match across spatial omics samples at single-cell resolution. Nature Methods 21, 1818–1829 (2024). 10.1038/s41592-024-02410-7

55 Yu, Y. & Xie, Z. Spatial Transcriptomic Alignment, Integration, and de novo 3D Reconstruction by STAIR. (2024). 10.21203/rs.3.rs-3939678/v1

56 Maynard, K. R. et al. Transcriptome-scale spatial gene expression in the human dorsolateral prefrontal cortex. Nat Neurosci 24, 425–436 (2021). 10.1038/s41593-020-00787-0

57 Zeng, H. et al. Integrative in situ mapping of single-cell transcriptional states and tissue histopathology in a mouse model of Alzheimer’s disease. Nat Neurosci 26, 430–446 (2023). 10.1038/s41593-022-01251-x

58 Tan, J. P. et al. A three-dimensional spatial transcriptome atlas reconstructs early organogenesis in primate Carnegie stages 9 and 10 embryos. Nat Cell Biol 28, 1309–1327 (2026). 10.1038/s41556-026-01956-2

59 Tian, R. et al. Integrating cross-sample and cross-modal data for spatial transcriptomics and metabolomics with SpatialMETA. Nat Commun 16, 8855 (2025). 10.1038/s41467-025-63915-z

60 Ren, P. et al. Systematic benchmarking of high-throughput subcellular spatial transcriptomics platforms across human tumors. Nat Commun 16, 9232 (2025). 10.1038/s41467-025-64292-3

61 Pratapa, A. et al. SAME: Topology-flexible transforms enable robust integration of multimodal spatial omics.bioRxiv, 2025.2007.2012.664419 (2025). 10.1101/2025.07.12.664419

62 Wang, H. et al. SPCoral: diagonal integration of spatial multi-omics across diverse modalities and technologies.bioRxiv, 2026.2002.2002.703207 (2026). 10.64898/2026.02.02.703207

63 Li, H., Zhang, Z., Squires, M., Chen, X. & Zhang, X. scMultiSim: simulation of single-cell multi-omics and spatial data guided by gene regulatory networks and cell-cell interactions. Nat Methods 22, 982–993 (2025). 10.1038/s41592-025-02651-0

64 Chen, P. et al. Integrating spatial omics and single-cell mass spectrometry imaging reveals tumor-host metabolic interplay in hepatocellular carcinoma. Proc Natl Acad Sci U S A 122, e2505789122 (2025). 10.1073/pnas.2505789122

65 Hu, J. et al. Multi-omic profiling of clear cell renal cell carcinoma identifies metabolic reprogramming associated with disease progression. Nat Genet 56, 442–457 (2024). 10.1038/s41588-024-01662-5

66 Lin-Shiao, E. et al. p63 establishes epithelial enhancers at critical craniofacial development genes. Sci Adv 5, eaaw0946 (2019). 10.1126/sciadv.aaw0946

67 Byrd, K. M. et al. Heterogeneity within Stratified Epithelial Stem Cell Populations Maintains the Oral Mucosa in Response to Physiological Stress. Cell Stem Cell 25, 814–829 e816 (2019). 10.1016/j.stem.2019.11.005

68 Blanpain, C. & Fuchs, E. Epidermal homeostasis: a balancing act of stem cells in the skin. Nat Rev Mol Cell Biol 10, 207–217 (2009). 10.1038/nrm2636

69 Cuylen, S. et al. Ki-67 acts as a biological surfactant to disperse mitotic chromosomes. Nature 535, 308–312 (2016). 10.1038/nature18610

70 Tumbar, T. et al. Defining the epithelial stem cell niche in skin. Science 303, 359–363 (2004). 10.1126/science.1092436

71 Vermeulen, M. et al. Quantitative interaction proteomics and genome-wide profiling of epigenetic histone marks and their readers. Cell 142, 967–980 (2010). 10.1016/j.cell.2010.08.020

72 Doupe, D. P. et al. A single progenitor population switches behavior to maintain and repair esophageal epithelium.Science 337, 1091–1093 (2012). 10.1126/science.1218835

73 Han, X. et al. Mapping the Mouse Cell Atlas by Microwell-Seq. Cell 173, 1307 (2018). 10.1016/j.cell.2018.05.012

74 Chen, L. et al. The short isoform of the CEACAM1 receptor in intestinal T cells regulates mucosal immunity and homeostasis via Tfh cell induction. Immunity 37, 930–946 (2012). 10.1016/j.immuni.2012.07.016

75 Williams, D. W. et al. Human oral mucosa cell atlas reveals a stromal-neutrophil axis regulating tissue immunity.Cell 184, 4090–4104 e4015 (2021). 10.1016/j.cell.2021.05.013

76 Garlanda, C., Dinarello, C. A. & Mantovani, A. The interleukin-1 family: back to the future. Immunity 39, 1003–1018 (2013). 10.1016/j.immuni.2013.11.010

77 Fu, X. et al. Spatial gene expression at single-cell resolution from histology using deep learning with GHIST. Nat Methods 22, 1900–1910 (2025). 10.1038/s41592-025-02795-z

78 Chen, W. et al. A visual-omics foundation model to bridge histopathology with spatial transcriptomics. Nat Methods 22, 1568–1582 (2025). 10.1038/s41592-025-02707-1

79 Hoang, D.-T. et al. A deep-learning framework to predict cancer treatment response from histopathology images through imputed transcriptomics. Nature Cancer 5, 1305–1317 (2024). 10.1038/s43018-024-00793-2

80 Zhang, D. et al. Inferring super-resolution tissue architecture by integrating spatial transcriptomics with histology.Nat Biotechnol 42, 1372–1377 (2024). 10.1038/s41587-023-02019-9

81 Min, W., Shi, Z., Zhang, J., Wan, J. & Wang, C. Multimodal contrastive learning for spatial gene expression prediction using histology images. Brief Bioinform 25 (2024). 10.1093/bib/bbae551

82 He, B. et al. Integrating spatial gene expression and breast tumour morphology via deep learning. Nat Biomed Eng 4, 827–834 (2020). 10.1038/s41551-020-0578-x

83 Qiu, X. et al. Spatiotemporal modeling of molecular holograms. Cell 187, 7351–7373.e7361 (2024). 10.1016/j.cell.2024.10.011

84 Dai, B. et al. A lightweight, ultrafast and general embedding framework for large-scale spatial omics data. bioRxiv,2026.2002.2004.703814 (2026). 10.64898/2026.02.04.703814

85 Lorensen, W. E. & Cline, H. E. in Seminal graphics: pioneering efforts that shaped the field 347–353 (1998).

86 Cao, J. et al. The single-cell transcriptional landscape of mammalian organogenesis. Nature 566, 496–502 (2019). 10.1038/s41586-019-0969-x

87 Sampath Kumar, A. et al. Spatiotemporal transcriptomic maps of whole mouse embryos at the onset of organogenesis. Nat Genet 55, 1176–1185 (2023). 10.1038/s41588-023-01435-6

88 Amorim, J. P. et al. A Conserved Notochord Enhancer Controls Pancreas Development in Vertebrates. Cell Rep32, 107862 (2020). 10.1016/j.celrep.2020.107862

89 Klar, A., Baldassare, M. & Jessell, T. M. F-spondin: a gene expressed at high levels in the floor plate encodes a secreted protein that promotes neural cell adhesion and neurite extension. Cell 69, 95–110 (1992). 10.1016/0092-8674(92)90121-r

90 Arekatla, G. et al. Identification of an embryonic differentiation stage marked by Sox1 and FoxA2 co-expression using combined cell tracking and high dimensional protein imaging. Nat Commun 15, 7860 (2024). 10.1038/s41467-024-52069-z

91 Delas, M. J. et al. Developmental cell fate choice in neural tube progenitors employs two distinct cis-regulatory strategies. Dev Cell 58, 3–17 e18 (2023). 10.1016/j.devcel.2022.11.016

92 Oh, S. W. et al. A mesoscale connectome of the mouse brain. Nature 508, 207–214 (2014). 10.1038/nature13186

93 Wang, Q. et al. The Allen Mouse Brain Common Coordinate Framework: A 3D Reference Atlas. Cell 181, 936–953 e920 (2020). 10.1016/j.cell.2020.04.007

94 Chen, X. et al. Whole-cortex in situ sequencing reveals input-dependent area identity. Nature 647, 203–212 (2025). 10.1038/s41586-024-07221-6

95 Han, L. et al. Single-cell spatial transcriptomic atlas of the whole mouse brain. Neuron 113, 2141–2160 e2149 (2025). 10.1016/j.neuron.2025.02.015

96 Shi, H. et al. Spatial atlas of the mouse central nervous system at molecular resolution. Nature 622, 552–561 (2023). 10.1038/s41586-023-06569-5

97 Yao, Z. et al. A high-resolution transcriptomic and spatial atlas of cell types in the whole mouse brain. Nature 624, 317–332 (2023). 10.1038/s41586-023-06812-z

98 Zhang, M. et al. Molecularly defined and spatially resolved cell atlas of the whole mouse brain. Nature 624, 343–354 (2023). 10.1038/s41586-023-06808-9

99 Lin, S. et al. Bridging the dimensional gap from planar spatial transcriptomics to 3D cell atlases. Nat Methods 23, 360–372 (2026). 10.1038/s41592-025-02969-9

100 Coleman, K., Schroeder, A. & Li, M. Unlocking the power of spatial omics with AI. Nat Methods 21, 1378–1381 (2024). 10.1038/s41592-024-02363-x

101 Liu, Y., Dai, Y. & Wang, L. Spatial omics at the forefront: emerging technologies, analytical innovations, and clinical applications. Cancer Cell 44, 24–49 (2026). 10.1016/j.ccell.2025.12.009

102 Skalli, O. et al. Alpha-smooth muscle actin, a differentiation marker of smooth muscle cells, is present in microfilamentous bundles of pericytes. J Histochem Cytochem 37, 315–321 (1989). 10.1177/37.3.2918221

103 Lertkiatmongkol, P., Liao, D., Mei, H., Hu, Y. & Newman, P. J. Endothelial functions of platelet/endothelial cell adhesion molecule-1 (CD31). Curr Opin Hematol 23, 253–259 (2016). 10.1097/moh.0000000000000239

104 Uhlar, C. M. & Whitehead, A. S. Serum amyloid A, the major vertebrate acute-phase reactant. European Journal of Biochemistry 265, 501–523 (1999). 10.1046/j.1432-1327.1999.00657.x

105 West, E. E., Woodruff, T., Fremeaux-Bacchi, V. & Kemper, C. Complement in human disease: approved and up-and-coming therapeutics. Lancet 403, 392–405 (2024). 10.1016/s0140-6736(23)01524-6

106 Yoshimura, A., Naka, T. & Kubo, M. SOCS proteins, cytokine signalling and immune regulation. Nature Reviews Immunology 7, 454–465 (2007). 10.1038/nri2093

107 Mantovani, A., Marchesi, F., Malesci, A., Laghi, L. & Allavena, P. Tumour-associated macrophages as treatment targets in oncology. Nature reviews Clinical oncology 14, 399–416 (2017).

108 Liu, Y. et al. Conserved spatial subtypes and cellular neighborhoods of cancer-associated fibroblasts revealed by single-cell spatial multi-omics. Cancer Cell 43, 905–924 e906 (2025). 10.1016/j.ccell.2025.03.004

109 Andrés-Sánchez, N., Fisher, D. & Krasinska, L. Physiological functions and roles in cancer of the proliferation marker Ki-67. J Cell Sci 135 (2022). 10.1242/jcs.258932

110 Upadhye, A. et al. Intra-tumoral T cells in pediatric brain tumors display clonal expansion and effector properties.Nat Cancer 5, 791–807 (2024). 10.1038/s43018-023-00706-9

111 Mariathasan, S. et al. TGFβ attenuates tumour response to PD-L1 blockade by contributing to exclusion of T cells.Nature 554, 544–548 (2018). 10.1038/nature25501

112 Iglesia, M. D. et al. Differential chromatin accessibility and transcriptional dynamics define breast cancer subtypes and their lineages. Nature Cancer 5, 1713–1736 (2024). 10.1038/s43018-024-00773-6

113 Rebuffet, L. et al. High-dimensional single-cell analysis of human natural killer cell heterogeneity. Nature Immunology 25, 1474–1488 (2024). 10.1038/s41590-024-01883-0

114 Chen, X. et al. An oncolytic virus delivering tumor-irrelevant bystander T cell epitopes induces anti-tumor immunity and potentiates cancer immunotherapy. Nature Cancer 5, 1063–1081 (2024). 10.1038/s43018-024-00760-x

115 Han, Y. et al. Spatiotemporal analyses of the pan-cancer single-cell landscape reveal widespread profibrotic ecotypes associated with tumor immunity. Nature Cancer 6, 1880–1898 (2025). 10.1038/s43018-025-01039-5

116 Zhang, M. S. et al. Hypoxia-induced macropinocytosis represents a metabolic route for liver cancer. Nature Communications 13, 954 (2022). 10.1038/s41467-022-28618-9

117 McDonald, P. C. et al. Regulation of pH by Carbonic Anhydrase 9 Mediates Survival of Pancreatic Cancer Cells With Activated KRAS in Response to Hypoxia. Gastroenterology 157, 823–837 (2019). 10.1053/j.gastro.2019.05.004

118 Peng, Q. et al. Dihydroartemisinin inhibited the Warburg effect through YAP1/SLC2A1 pathway in hepatocellular carcinoma. J Nat Med 77, 28–40 (2023). 10.1007/s11418-022-01641-2

119 Shen, H. et al. Lactate Metabolic Reprogramming Mediated by CircRNA-LDHA Complex Facilitates Innate Immune Evasion of Liver Cancer. Adv Sci (Weinh) 12, e09989 (2025). 10.1002/advs.202509989

120 Tauriello, D. V. F. et al. TGFβ drives immune evasion in genetically reconstituted colon cancer metastasis. Nature554, 538–543 (2018). 10.1038/nature25492

121 LaGory, E. L. & Giaccia, A. J. The ever-expanding role of HIF in tumour and stromal biology. Nat Cell Biol 18, 356–365 (2016). 10.1038/ncb3330

122 Derynck, R. & Zhang, Y. E. Smad-dependent and Smad-independent pathways in TGF-beta family signalling.Nature 425, 577–584 (2003). 10.1038/nature02006

123 David, C. J. & Massagué, J. Contextual determinants of TGFβ action in development, immunity and cancer. Nature Reviews Molecular Cell Biology 19, 419–435 (2018). 10.1038/s41580-018-0007-0

124 Zhang, Y. E. Non-Smad pathways in TGF-β signaling. Cell Research 19, 128–139 (2009). 10.1038/cr.2008.328

125 Shi, M. et al. Latent TGF-β structure and activation. Nature 474, 343–349 (2011).10.1038/nature10152

126 Barrow, H. G., Tenenbaum, J. M., Bolles, R. C. & Wolf, H. C. in International Joint Conference on Artificial Intelligence.

127 Klein, D. et al. Mapping cells through time and space with moscot. Nature 638, 1065–1075 (2025). 10.1038/s41586-024-08453-2

128 He, G., Xiao, Y., Xu, Z., Zhou, X. & Peng, S. ERNet: Efficient Non-Rigid Registration Network for Point Sequences. 2025 IEEE/CVF International Conference on Computer Vision (ICCV), 27156–27165 (2025).

129 Yew, Z. J. & Lee, G. H. RPM-Net: Robust Point Matching Using Learned Features. 2020 IEEE/CVF Conference on Computer Vision and Pattern Recognition (CVPR), 11821–11830 (2020).

130 Bao, H., Dong, L., Piao, S. & Wei, F. Beit: Bert pre-training of image transformers. arXiv preprint arXiv:2106.08254 (2021).

131 Vaswani, A. et al. Attention is all you need. Advances in neural information processing systems 30 (2017).

132 Xiong, R. et al. in International conference on machine learning. 10524–10533 (PMLR).

133 Ganin, Y. & Lempitsky, V. in International conference on machine learning. 1180–1189 (PMLR).

134 Virshup, I., Rybakov, S., Theis, F. J., Angerer, P. & Wolf, F. A. anndata: Access and store annotated data matrices.Journal of Open Source Software 9, 4371 (2024).

135 Stringer, C. & Pachitariu, M. Cellpose3: one-click image restoration for improved cellular segmentation. Nat Methods 22, 592–599 (2025). 10.1038/s41592-025-02595-5

136 Wolf, F. A., Angerer, P. & Theis, F. J. SCANPY: large-scale single-cell gene expression data analysis. Genome Biol 19, 15 (2018). 10.1186/s13059-017-1382-0

137 Chen, R. J. et al. Towards a general-purpose foundation model for computational pathology. Nature Medicine 30, 850–862 (2024). 10.1038/s41591-024-02857-3

138 Lorensen, W. E. & Cline, H. E. Marching cubes: A high resolution 3D surface construction algorithm. SIGGRAPH Comput. Graph. 21, 163–169 (1987). 10.1145/37402.37422

139 Pedregosa, F. et al. Scikit-learn: Machine learning in Python. the Journal of machine Learning research 12, 2825–2830 (2011).

140 Varrone, M., Tavernari, D., Santamaria-Martinez, A., Walsh, L. A. & Ciriello, G. CellCharter reveals spatial cell niches associated with tissue remodeling and cell plasticity. Nat Genet 56, 74–84 (2024).

